# ER-mitochondria distance is a critical parameter for efficient mitochondrial Ca^2+^ uptake and oxidative metabolism

**DOI:** 10.1101/2024.07.24.604907

**Authors:** Giulia Dematteis, Laura Tapella, Claudio Casali, Maria Talmon, Elisa Tonelli, Simone Reano, Adele Ariotti, Emanuela Pessolano, Justyna Malecka, Gabriela Chrostek, Gabrielė Kulkovienė, Danielius Umbrasas, Carla Distasi, Mariagrazia Grilli, Graham Ladds, Nicoletta Filigheddu, Luigia G Fresu, Katsuhiko Mikoshiba, Carlos Matute, Paula Ramos-Gonzalez, Aiste Jekabsone, Tito Calì, Marisa Brini, Marco Biggiogera, Fabio Cavaliere, Riccardo Miggiano, Armando A Genazzani, Dmitry Lim

**Author notes:** Department of Drug Science and Technology, University of Turin, Turin, Italy (Armando A Genazzani).

## Abstract

IP_3_ receptor (IP_3_R)-mediated Ca^2+^ transfer at the mitochondria-endoplasmic reticulum (ER) contact sites (MERCS) drives mitochondrial Ca^2+^ uptake and oxidative metabolism and is linked to different pathologies, including Parkinson’s disease (PD). The dependence of Ca^2+^ transfer efficiency on the ER-mitochondria distance remains unexplored. Employing molecular rulers that stabilize ER-mitochondrial distances at 5 nm resolution, and using genetically-encoded Ca^2+^ indicators targeting the ER lumen and the sub-mitochondrial compartments, we now show that a distance of ∼20 nm is optimal for Ca^2+^ transfer and mitochondrial oxidative metabolism due to enrichment of IP_3_R at MERCS. In human iPSC-derived astrocytes from PD patients, 20 nm MERCS were specifically reduced which correlated with a reduction of mitochondrial Ca^2+^ uptake. Our work determines with precision the optimal distance for Ca^2+^ flux between ER and mitochondria and suggests a new paradigm for fine control over mitochondrial function.

---

Ca^2+^ signals are required to drive mitochondrial bioenergetics through the activation of enzymes in the mitochondrial matrix ^1,2^ and deficiency or excessive mitochondrial Ca^2+^ signals have been associated with cellular dysfunction and disease pathogenesis ^3–6^. Mitochondria take up Ca^2+^ through a low-affinity mitochondrial Ca^2+^ uniporter complex (mtCU) ^7^. Close apposition of the endoplasmic reticulum (ER) membrane and the outer mitochondrial membrane (OMM) at the mitochondria-ER contact sites (MERCS) leads to high [Ca^2+^] hot spots and a direct transfer of Ca^2+^ through the ER-located inositol-1,4,5-trisphosphate receptors (IP_3_Rs) and OMM-located porin/voltage-dependent cation channel 1 (VDAC1) ^8–11^. The distance between the ER and the OMM, at which Ca^2+^ transfer from the ER to the mitochondrial matrix occurs, has been proposed to lay within the 10-25 nm range ^12–15^. However, it is not known whether ER-mitochondrial Ca^2+^ transfer operates with the same efficiency in a range of ER-OMM distances or if a specific distance exists for optimal Ca^2+^ flux through IP_3_R to mitochondria (Fig. 1a).

**Fig. 1.**
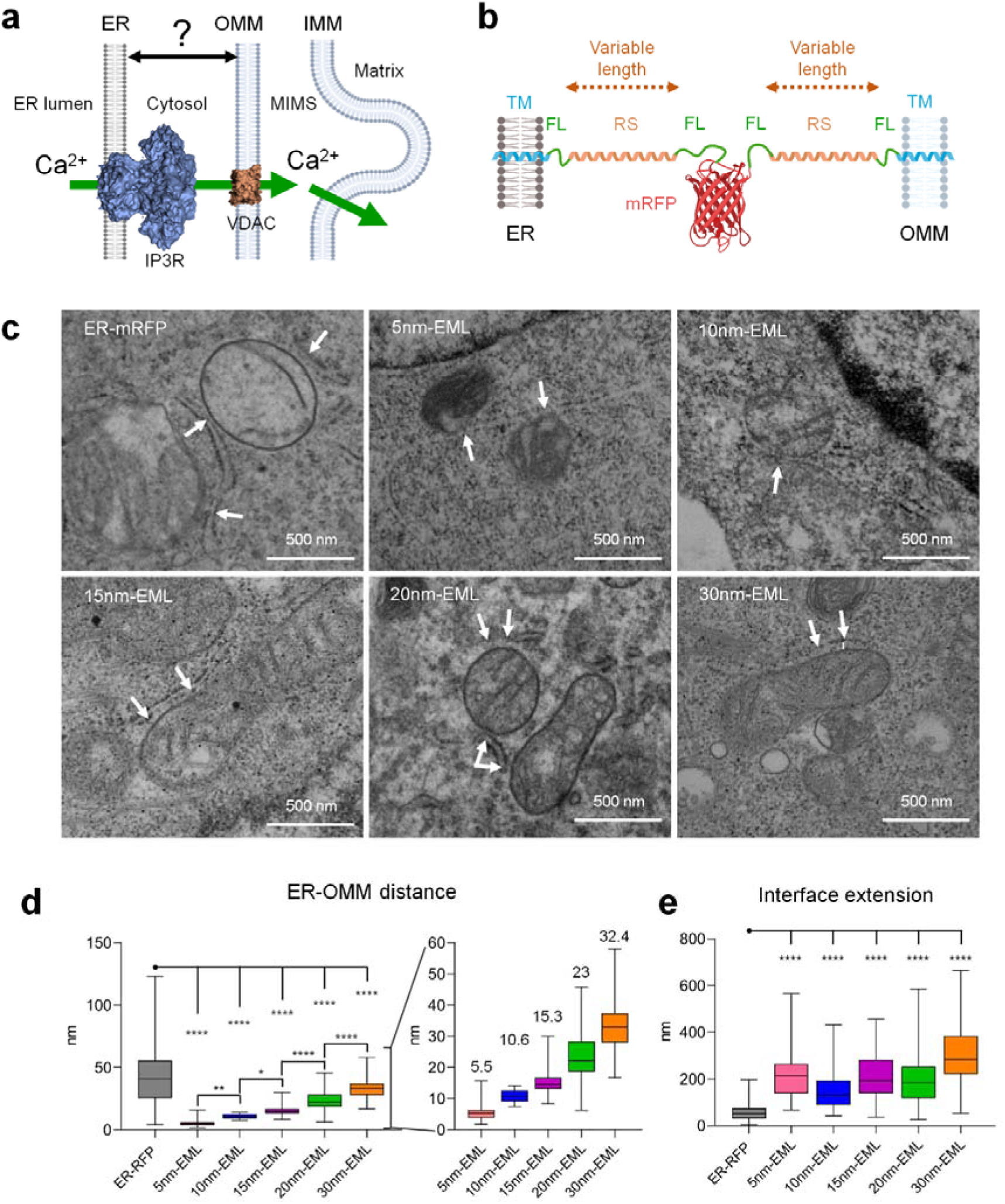
Characterization of ER-mitochondrial linkers (EMLs). (**a**) A scheme depicting question addressed in this study: what is the distance between ER membrane and OMM for optimal Ca^2+^ flux? Lipid bilayer, IP_3_R and VDAC are represented in scale. (**b**) Scheme illustrating design of ER-mitochondrial linkers. TM, transmembrane domain; FL, flexible linker; RS, rigid spacer; ER, endoplasmic reticulum; OMM, outer mitochondrial membrane; mRFP, monomeric red fluorescent protein. (**c**) TEM validation of EMLs overexpressed in HeLa cells. Arrows indicate ER membrane adjacent to mitochondria. Scale bar = 500 nm. (**d**) Quantification of ER-OMM distance on TEM images. Inset scales up quantified distances and shows the average distance per EML. (**e**) Quantification of the length of the interface between ER and mitochondria. One-way ANOVA, Tukey post hoc test, n = 31-178 contacts from at least 15 cells per condition. * p < 0.05, ** p < 0.01, **** p < 0.0001.

To address this question, we employed a palette of existent and newly generated synthetic ER-mitochondrial linkers (EML) designed to maintain the distance between ER and OMM in a range from 5 to 30 nm with a spatial resolution of 5 nm and genetically-encoded Ca^2+^ indicators targeting the ER lumen or the sub-mitochondrial compartments.

## RESULTS

### Generation of the extended palette of ER-mitochondrial linkers

We used five EMLs fixing ER-OMM distance at ≤5 nm (denominated as 5nm-EML), ≤10-12nm (10nm-EML), 15nm, 20nm and 30nm (15nm-EML, 20nm-EML and 30nm-EML, respectively). 5nm-EML and 10nm-EML, kindly provided by Gyorgy Hajnoczky, Thomas Jefferson University, were published elsewhere ^12,16^. 15nm-EML, 20nm-EML, and 30nm-EML were generated *de novo* (Fig. 1b, Extended Data Fig. 1) and were composed of monomeric red fluorescent protein (mRFP) flanked by a rigid α-helical spacer of a defined length, derived from the structure of myosin-VI known for its exceptional rigidity ^17^ (PDB code: 6OBI). Structural prediction, using Robetta service ^18^ was used to select the correct number of amino acids resulting in the desired ruler lengths. At the N- and C-termini, targeting sequences to OMM (AKAP1) and to the ER membrane (Ubc6) ^16^, were attached, respectively (Extended Data Fig. 1). All linkers were validated by electron microscopy (Fig. 1c), and their ability to impose the defined distance has been confirmed (Fig. 1d). Moreover, all linkers significantly increased the length of the interaction between the two organelles (Fig. 1e). Importantly, none of the EMLs affected cell viability up to 72 h post-transfection (Extended Data Fig. 2a). Similarly, no changes of total levels of proteins, implicated in mitochondrial dynamics, such as dynamin-related protein (DRP1), p-DRP1, mitofusin 1 (MFN1) and MFN2 ^19^ (Extended Data Fig. 2b) or in Ca^2+^ uptake (mitochondrial Ca^2+^ uniporter, MCU) ^20,21^ (Extended Data Fig. 2c) were observed. Localization of EMLs was confirmed using confocal microscopy (Extended Data Fig. 3). These data demonstrate that EML overexpression in HeLa cells is a suitable model to study the correlation between ER-mitochondrial distance and Ca^2+^ transfer.

### A bell-shaped ER-OMM distance-dependence of the ER-mitochondrial Ca^2+^ flux with a maximum at ∼**20 nm.**

Assessment of mitochondrial calcium uptake, using 4mtD3cpv probe targeted to the mitochondrial matrix ^22^ 48 h after co-transfection with EMLs, revealed that the amplitude of ATP-induced Ca^2+^ transients in the mitochondrial matrix ([Ca^2+^]_M_) in 5nm- and 10nm-EML-expressing cells was significantly reduced compared with control cells expressing ER-mRFP (Fig. 2a). Overexpression of 15nm-EML had no effect, while 20nm-EML strongly enhanced ATP-evoked [Ca^2+^]_M_ transient. Overexpression of 30nm-EML resulted in a drastic reduction of ATP-evoked [Ca^2+^]_M_ signals (Fig. 2a, b). These results confirm previous observations and predictions that when the ER-OMM distance is too short (5nm-EML) or too long (30nm-EML) the Ca^2+^ transfer is inefficient ^14–16^. Intriguingly, 10nm-EML strongly suppressed, while 15nm-EML did not change ER-mitochondrial Ca^2+^ flux compared with control cells (Fig. 2a, b) in spite of significant increase of the interface length between the membranes (Fig. 1e).

**Fig. 2.**
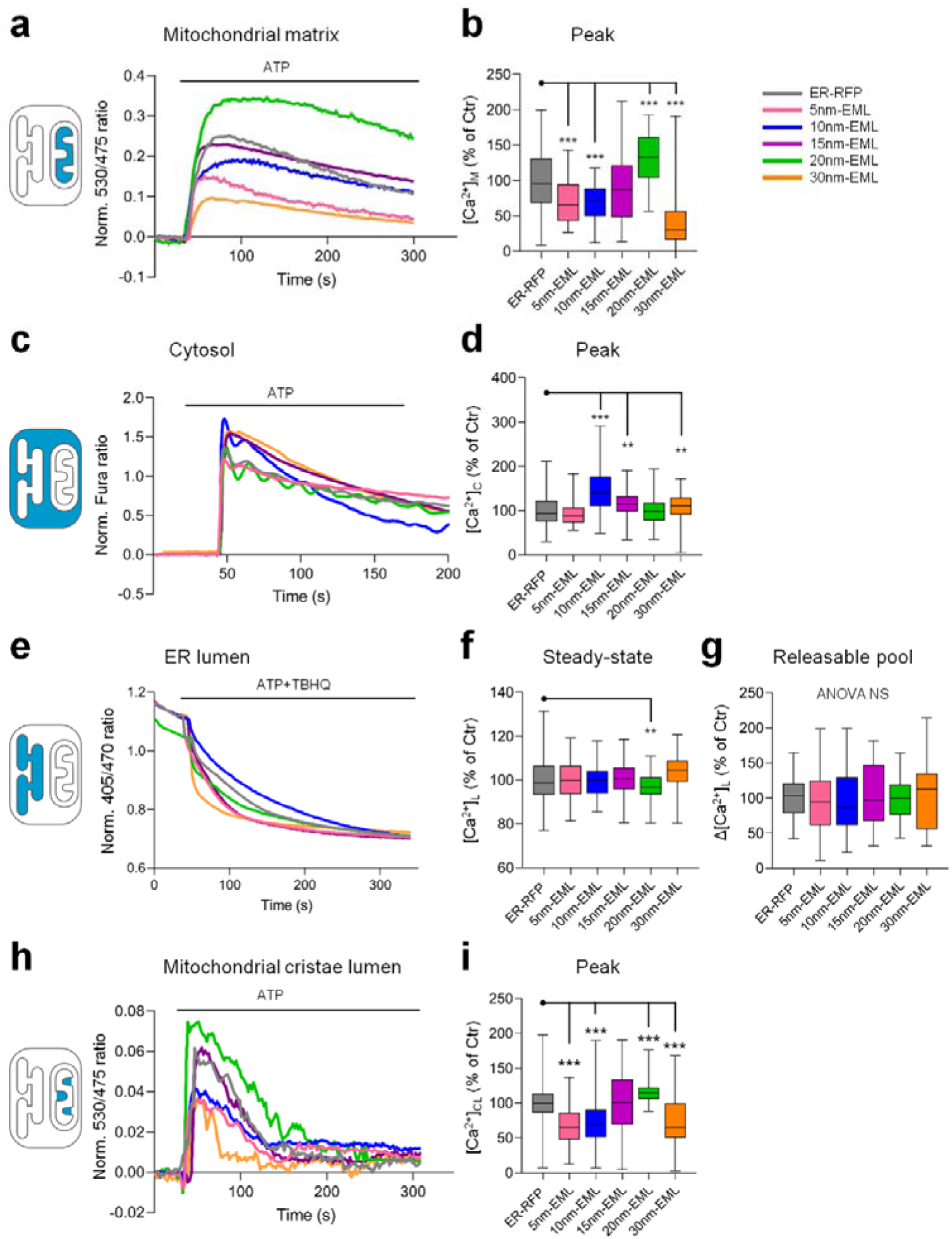
Effects of EMLs on cellular Ca^2+^ homeostasis and ER-mitochondrial Ca^2+^ transfer. Representative traces and quantifications of Ca^2+^ signals in the mitochondrial matrix (**a, b**), cytosol (**c, d**), ER lumen, (**e, f and g**) and mitochondrial cristae lumen (**h, i**). Whisker plots of data collected from 88-150 (**b**), 130-492 (**d**), 91-260 (**f, g**), 75-212 (**i**) cells from at least 3 independent coverslips analyzed from 3 independent experiments. One-way ANOVA, Tukey post hoc test. ** p < 0.01, ***, p < 0.001.

To investigate if alterations of Ca^2+^ handling in cytosol or ER, upon EMLs overexpression, could affect [Ca^2+^]_M_ response, we first assessed ATP-induced IP_3_R-mediated Ca^2+^ dynamics in the cytosol ([Ca^2+^]_C_) using Fura-2 probe ^23^. As shown in Fig. 2c and d, in 5nm- and 20nm-EML-expressing cells there were no differences in ATP-evoked [Ca^2+^]_C_ transient. In 10nm-EML, 15nm- and 30nm-EML-expressing cells [Ca^2+^]_C_ response was enhanced. Resting steady-state luminal ER Ca^2+^ levels ([Ca^2+^]_L_) and the ER releasable Ca^2+^ pool (Δ[Ca^2+^]_L_) were measured using a green fluorescent protein (GFP)-aequorin fusion protein (GAP3) probe ^24^. The GAP3-transduced HeLa cells were stimulated with a cocktail, containing ATP (100 µM) and tert-butyl hydroquinone (TBHQ, 100 µM) in a Ca^2+^-free KRB solution supplemented with 500 µM EGTA, to induce rapid and complete ER depletion (Fig. 2e). No significant differences in [Ca^2+^]_L_ and Δ[Ca^2+^]_L_ were observed upon EML overexpression, with the exception of 20nm-EML, which resulted in a modest but significant reduction of the steady-state [Ca^2+^]_L_ compared with control cells expressing ER-mRFP (Fig. 2f). Nevertheless, the Δ[Ca^2+^]_L_ in 20nm-EML-expressing HeLa was not different from control (Fig. 2g). These results suggest that the changes of ATP-evoked [Ca^2+^]_M_ transients in EMLs-expressing cells were not due to alterations of ER Ca^2+^ content and/or Ca^2+^ release capacity.

Potentially, alterations in mtCU activity could influence Ca^2+^ transients in the mitochondrial matrix. To confirm that the observed effects of EMLs (Fig. 2a) were independent of mtCU, we measured Ca^2+^ in the mitochondrial cristae lumen (CL) ([Ca^2+^]_CL_). Employing a recently developed ratiometric probe exploiting the targeting sequence from Reactive Oxygen Species Modulator 1 protein (denominated as ROMO-GemGeCO) ^25^, we show that, quantitatively, [Ca^2+^]_CL_ signals detected by ROMO-GemGeCO (Fig. 2h, i) faithfully resemble those of [Ca^2+^]_M_ measured by 4mtD3cpv (see Fig. 2a).

We also checked if the 20 nm ER-mitochondrial distance enhances Ca^2+^ transfer in other cellular types. We found that ATP-induced [Ca^2+^]_M_ was significantly potentiated in hepatocellular carcinoma Huh-7 cell line (Extended Data Fig. 4) and in primary murine embryonic fibroblasts (MEFs) transduced with 20nm-EML (see below). These results demonstrate that i) EMLs, based on rigid α-helical structures, represent a unique tool for the control of ER-OMM distance, allowing discrimination of events with ER-OMM distance resolution of 5 nm; ii) ER-mitochondrial Ca^2+^ flux critically depends on transversal ER-OMM distance and iii) the dependence is bell-shaped with ∼20 nm representing an optimal distance for both ER-mitochondria Ca^2+^ transfer and mitochondrial Ca^2+^ uptake.

### Enrichment of IP_3_R in 20 nm ER-OMM space with formation of functional IP_3_R-VDAC Ca^2+^ transferring units

To investigate if IP_3_R and VDAC were implicated in the increase of Ca^2+^ transfer in 20nm-EML-expressing cells ^26^ we used a number of complementary approaches focusing on 5nm-, 10nm- and 20nm-EMLs because they produced clear opposite effects on ER-mitochondrial Ca^2+^ flux. First, we performed a proximity ligation assay (PLA), widely exploited to study IP_3_R-VDAC1 juxtaposition (Fig. 3a). As shown in Fig. 3b, in control ER-mRFP-expressing cells, PLA signals show diffuse dotty patterns corresponding to juxtaposed (≤40nm) IP_3_R and VDAC1/3 proteins using anti-pan-IP_3_R and anti-VDAC1/3 antibodies, detecting putative close apposition of IP_3_R and VDAC1/3 at MERCS. Overexpression of 5nm-EML suppressed PLA signal, suggesting that at this distance the juxtaposition of IP_3_R and VDAC1/3 is inhibited. Overexpression of 10nm-EML resulted in a significant increase of PLA signal suggesting an enhanced juxtaposition of IP_3_R and VDAC1/3. However, a higher magnification examination shows only partial co-localization of PLA signal with 10nm-EML. Considering the reduced ATP-induced [Ca^2+^]_M_ and [Ca^2+^]_CL_ transients (Fig. 2), this result indicates that, although juxtaposed, IP_3_R and VDAC1/3 are unable to form functional Ca^2+^-transferring complexes. Strikingly, overexpression of 20nm-EML resulted in a strong increase of PLA signal (Fig. 3c) with a significantly higher co-localization between PLA signal and 20nm-EML (Fig. 3d), suggesting an enrichment of 20 nm MERCS with functional IP_3_R-VDAC complexes. Next, we performed immunofluorescent (IF) analysis using anti-pan-IP_3_R antibody, recognizing all three IP_3_R isoforms, which, in HeLa cells, form hetero-tetrameric IP_3_Rs ^27^, to analyze the distribution of IP_3_Rs in EMLs-expressing cells (Fig. 3e). Overexpression of 5nm-EML did not significantly alter the wide-spread distribution of IP_3_R in the cell, while 10nm- and 20nm-EMLs produced an enrichment of IP_3_Rs in the proximity of EMLs (Fig. 3f). In line with PLA data, Pearson correlation coefficient quantification showed a significantly higher co-localization between IP_3_R and 20nm-EML compared with 10nm-EML (Fig. 3g).

**Fig. 3.**
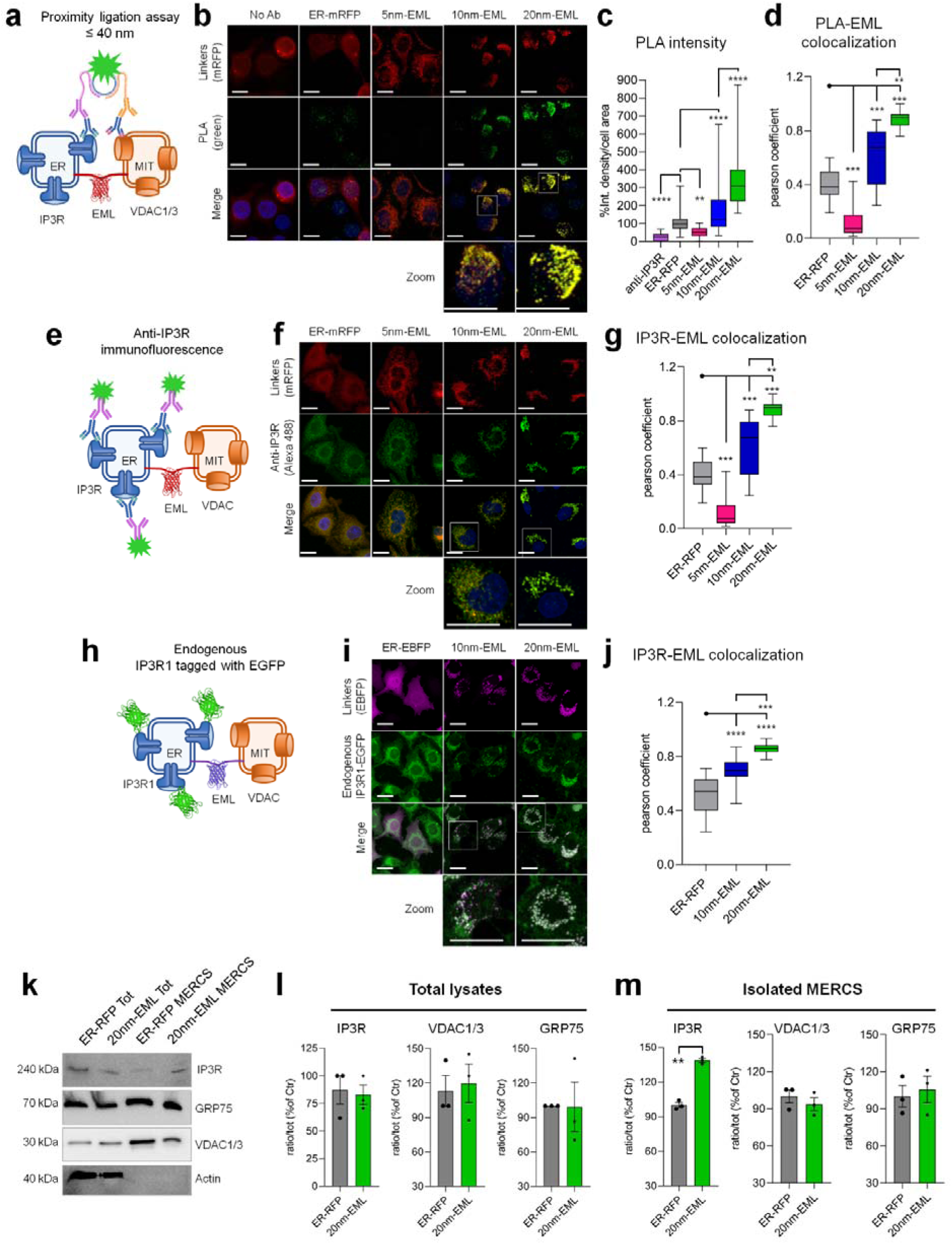
Effects of EMLs on localization of IP_3_R. (**a-d**) Proximity ligation assay (PLA) using anti-pan-IP_3_R, and anti-VDAC1/3 antibody (green) of HeLa cells expressing control plasmid (ER-mRFP) and ER-mitochondrial linkers (EMLs) (red). (**a**) Cartoon illustrating that only IP_3_Rs and VDAC1/3 juxta-positioned at ≤ 40 nm are labelled. Representative images (**b**) and quantification of the PLA-labelled area intensity of and PLA-EML Pearson colocalization coefficient (**d**). Scale bars = 20 µM. (**e-g**) Immunofluorescent analysis using anti-pan-IP_3_R antibody (green) in ER-mRFP and EML-expressing HeLa cells (red). (**e**) Cartoon illustrating that all IP_3_Rs present in the cells are labelled. Representative images (**f**) and quantification of IP_3_R-EML Pearson colocalization coefficient (**g**). Scale bars = 20 µM. (**h-j**) Localization of endogenous IP_3_R1 tagged with EGFP in HeLa cells (endogenous IP_3_R1-EGFP, green) upon expression of ER-EBFP, and 10nm- and 20nm-EMLs containing EBFP (magenta). (**h**) Cartoon illustrating that all endogenous IP_3_R1 receptors are tagged with EGFP. Representative images (**i**) and quantification of EGFP-EML Pearson colocalization coefficient (**j**). Scale bars = 20 µM. Whisker plots report data collected from 54-106 (**c**), 25-30 (**d**), 8-18 (**g**), 16-18 (**j**) cells from at least 3 independent coverslips analyzed from 3 independent cultures. One-way ANOVA, Tukey post hoc test. ** p < 0.01, *** p < 0.001, **** p < 0.0001. Representative images (**K**) and quantification of total cell lysates (**l**) and isolated MERCS (**m**) using anti-pan-IP_3_R antibody, anti-VDAC1/3 or anti-GRP75 antibody. Data are expressed as mean ± SEM of n = 3 independent experiments. Unpaired two-tailed Student’s t-test. **, p < 0.01.

Enrichment of IP_3_R fluorescent signal in the EML region and its strong reduction in the resting part of the cytosol in 20nm-EML expressing cells suggest relocation of IP_3_R into 20 nm ER-mitochondrial space. To confirm this hypothesis in an unbiased manner, independent of the enhancement of fluorescent signal during immunodecoration with first and second antibodies, we expressed 10nm- and 20nm-EMLs in a recently generated HeLa cell line, in which endogenous IP_3_R1 was tagged with EGFP using CRISPR/Cas9 technology (HeLa-IP_3_R1-EGFP) ^27^ (Fig. 3h). To avoid minimal possible contamination of EGFP signal by fluorescence deriving from mRFP-containing linker variants, we generated and validated EMLs expressing enhanced blue fluorescent protein (EBFP) (Extended Data Fig. 5). Overexpression of the control (ER-EBFP) construct in HeLa-IP_3_R1-EGFP cells did not alter the distribution of endogenous IP_3_R1-EGFP signal (Fig. 3i, ER-EBFP). However, overexpression of 10nm- and 20nm-EML-EBFP constructs, 48 h after transfection, resulted in strong re-localization of fluorescent signals in the area of EMLs. In line with PLA and IF analyses, co-localization between IP_3_R1-EGFP and EMLs was significantly higher for 20nm-EML compared with 10nm-EML (Fig. 3j). A residual signal remained in some intracellular areas or in correspondence to the edge of the cell. Western blot analysis of total lysates showed that the enrichment of IP_3_R in 20nm MERCS was not due to altered expression of IP_3_Rs (Fig. 3k, l and Extended Data Fig. 6a). Furthermore, in confirmation of the IP_3_R re-location hypothesis, IP_3_R protein was significantly increased in MERCS isolated from HeLa cells stably expressing 20nm-EML compared with cells expressing control ER-mRFP construct (Fig. 3k and m). Interestingly, the amount of VDAC1/3 or GRP75 proteins was not different from control cells either in total cell lysates (Fig. 3l and Extended Data Fig. 6b) or in MERCS isolated from 20nm-EML-overexpressing cells (Fig. 3m) suggesting that enrichment of IP_3_Rs in 20 nm MERCS was independent of VDAC1/3 localization.

The above-described experiments are based on a generally-accepted paradigm that both IP_3_R and VDAC are required for ER-mitochondrial Ca^2+^ transfer ^28,29^. However, forced membrane tethering could potentially predispose to an IP_3_R-independent Ca^2+^ release, e.g., via Ca^2+^ leak from the ER and/or activation of Ca^2+^-induced Ca^2+^ release mechanism ^30,31^. On the other hand, it has been suggested that several β-barrel-forming OMM proteins can be permeable to ions ^32,33^. To investigate whether IP_3_R and VDAC1/3 are the only ones responsible for the enhanced Ca^2+^ flux in 20nm-EML-expressing cells, we used HeLa cells with triple IP_3_R1,2,3 knock-out (HeLa-TKO) ^29^ and MEFs derived from VDAC1/3-KO mice ^34^. ATP-induced Ca^2+^ signals were monitored in mitochondrial CL using ROMO-GemGeCO probe. [Ca_2+_]_CL_ signals were not detected in ER-mRFP-expressing HeLa-TKO cells, nor in cells overexpressing either 10nm- or 20nm-EML (Extended Data Fig. 7a,b). Similarly, [Ca_2+_]_CL_ signals were completely absent in ATP-stimulated VDAC1/3- KO MEFs in either condition (Extended Data Fig. 7c,d). Notably, in both WT cells, HeLa and MEFs, [Ca_2+_]_CL_ signals were detected, and, in line with that reported in Fig. 2, 10nm-EML reduced, while 20nm-EML significantly enhanced Ca^2+^ transients in CL. Therefore, both IP_3_R and VDAC1/3 are necessary for Ca^2+^ transfer at 20 nm MERCS.

### 10 nm ER-OMM distance inhibits, while 20 nm distance promotes mitochondrial respiration and ATP production

One of the most accredited functions of Ca^2+^ signals in mitochondria is the regulation of mitochondrial energetics through the activation of metabolic enzymes and ATP synthase ^2^. Therefore, we investigated the effect of EMLs overexpression on mitochondrial respiration and ATP production using Oroboros oxygraphy (Fig. 4a). We found that 10nm-EML overexpression reduced, while the 20nm-EML one significantly increased both basal oxygen consumption (Fig. 4b), respiratory reserve (Fig. 4c) and ATP-linked respiration (Fig. 4d). These data were corroborated by quantification of total cellular ATP content (Fig. 4e). Potentially, variations of the mitochondrial membrane potential (ΔΨm), due to expression of EMLs, could influence respiration and ATP production. However, measurement of ΔΨm with JC-1 probe using flow cytometry, in cells expressing EBFP variant EMLs, did not reveal any difference in ΔΨm among ER-EBFP, 10nm-EML and 20nm-EML-expressing HeLa cells (Extended Data Fig. 8). Our results suggest that mitochondrial oxidative metabolism and ATP supply may be efficiently regulated, through regulation of Ca^2+^ flux, by modulating the distance between ER and mitochondria.

**Fig. 4.**
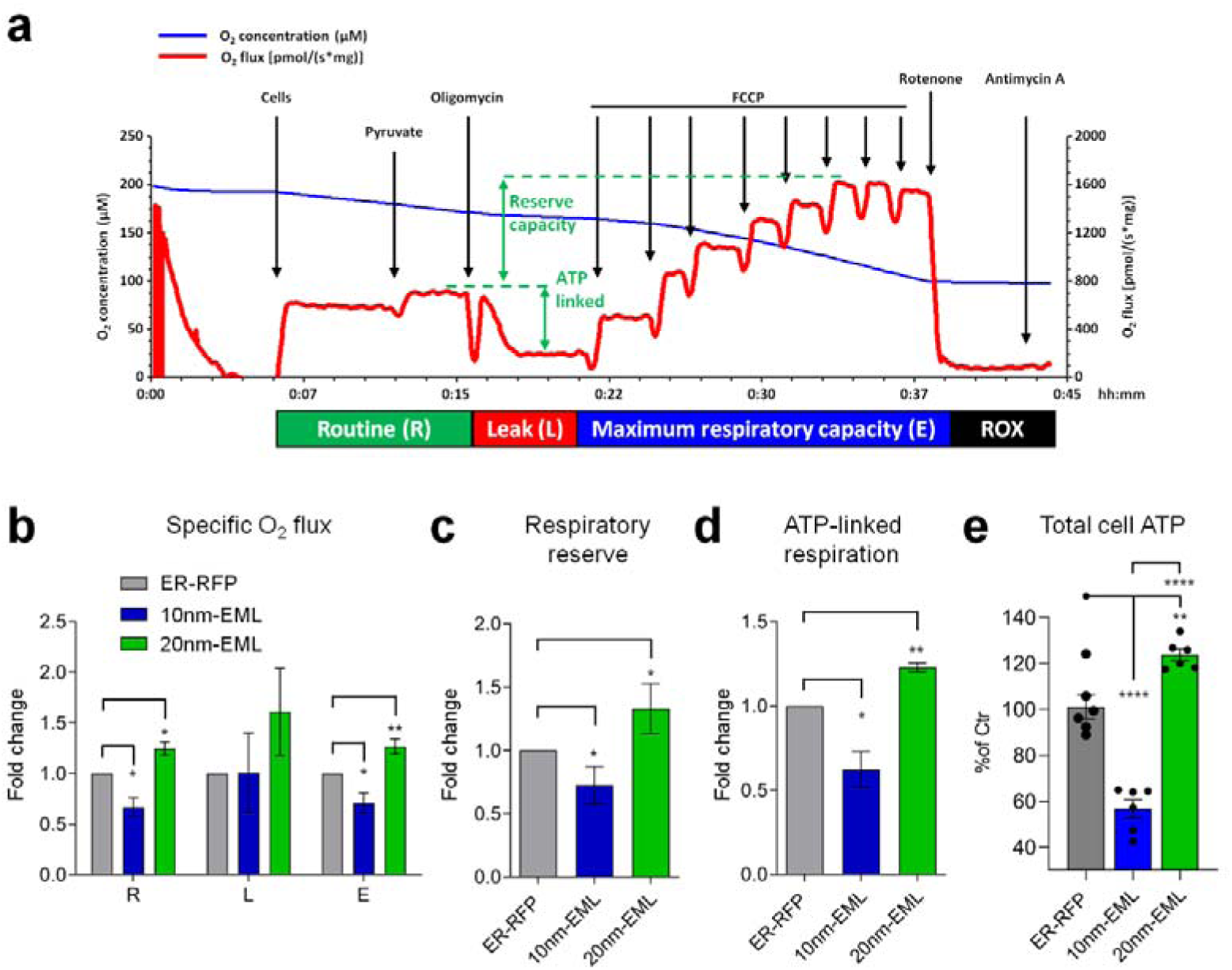
ER-OMM distance-dependent mitochondrial respiration and ATP production. (**a**). Representative traces (ER-RFP), illustrating the protocol of high-resolution respirometry for quantification of respiration in intact cells; blue line: oxygen concentration, red line: oxygen flux. (**b**) Oxygen flux in the routine state (R); in the leakage state (L) after addition of oligomycin, an inhibitor of ATP synthase; after the addition of FCCP, an uncoupler of oxidative phosphorylation to induce maximum respiratory capacity (E). All data are expressed as specific flux, i.e., oxygen consumption normalized to the sample protein content and after subtraction of non-mitochondrial oxygen flux (ROX). (**c**) Reserve respiratory capacity obtained by the subtraction of R from E. (**d**) Oxygen consumption linked to ATP production, i.e., oligomycin-sensitive respiration obtained by the subtraction of L from R. Data are expressed as mean ± SEM of fold changes above the control of n = 4 (ER-RFP and 10nm-EML) or n = 3 (ER-RFP and 20nm-EML) independent experiments. Unpaired two-tailed t-test 10nm-EML or 20nm-EML vs ER-RFP. * p-value < 0.05; ** p-value < 0.01. (**e**) Total cellular ATP. Data are expressed as mean ± SEM of % of control (ER-RFP) of n = 6 independent experiments. One-way ANOVA, Tukey post hoc test, ** p-value < 0.01; **** p-value < 0.0001.

### MERCS are disrupted in Parkinson’s disease astrocytes: rescue by overexpression of 20nm-EML

The data, obtained using overexpression of artificial ER-OMM tethers, suggest that 20 nm distance specifically promotes Ca^2+^ transfer between ER and mitochondria through preferential localization of IP_3_Rs at 20 nm MERCS. Estimation of physiological abundance of MERCS on TEM images suggests that 18-22 nm MERCS account for ∼18% of all MERCS ranging from 5 to 80 nm, indicating that almost 1/5 of MERCS are potentially deputed to Ca^2+^ transfer, although this percentage likely depends on cell type and condition ^14^.

Increasing evidence suggests that MERCS are altered in neurodegeneration, including Alzheimer’s (AD) and Parkinson’s (PD) diseases ^15,35–37^. To investigate whether specific alteration of 20 nm MERCS may mediate mitochondrial Ca^2+^ deficiency in pathological conditions, we took advantage of a recently reported split-GFP contact site sensors (SPLICS) ^38,39^, which we adapted to reconstitute bright GFP fluorescence specifically at 20 nm between ER and OMM (Fig. 5a and Extended Data Fig. 9). Transfection of 20nm-SPLICS resulted in appearance of bright fluorescent dots distributed throughout the cell in sites of the higher ER and mitochondrial density (Extended Data Fig. 10a). Co-expression of 20nm-SPLICS probe with 20nm-EML resulted in a drastic increase of SPLICS signal and its complete co-localization with 20nm-EML (Extended Data Fig. 10b), confirming proper functioning of the probe, while immunodecoration of IP_3_Rs showed close juxtaposition of 20nm-SPLICS with a fraction of IP_3_Rs, although, expectedly, SPLICS-free IP_3_Rs were also detected (Extended Data Fig. 10c).

**Fig. 5.**
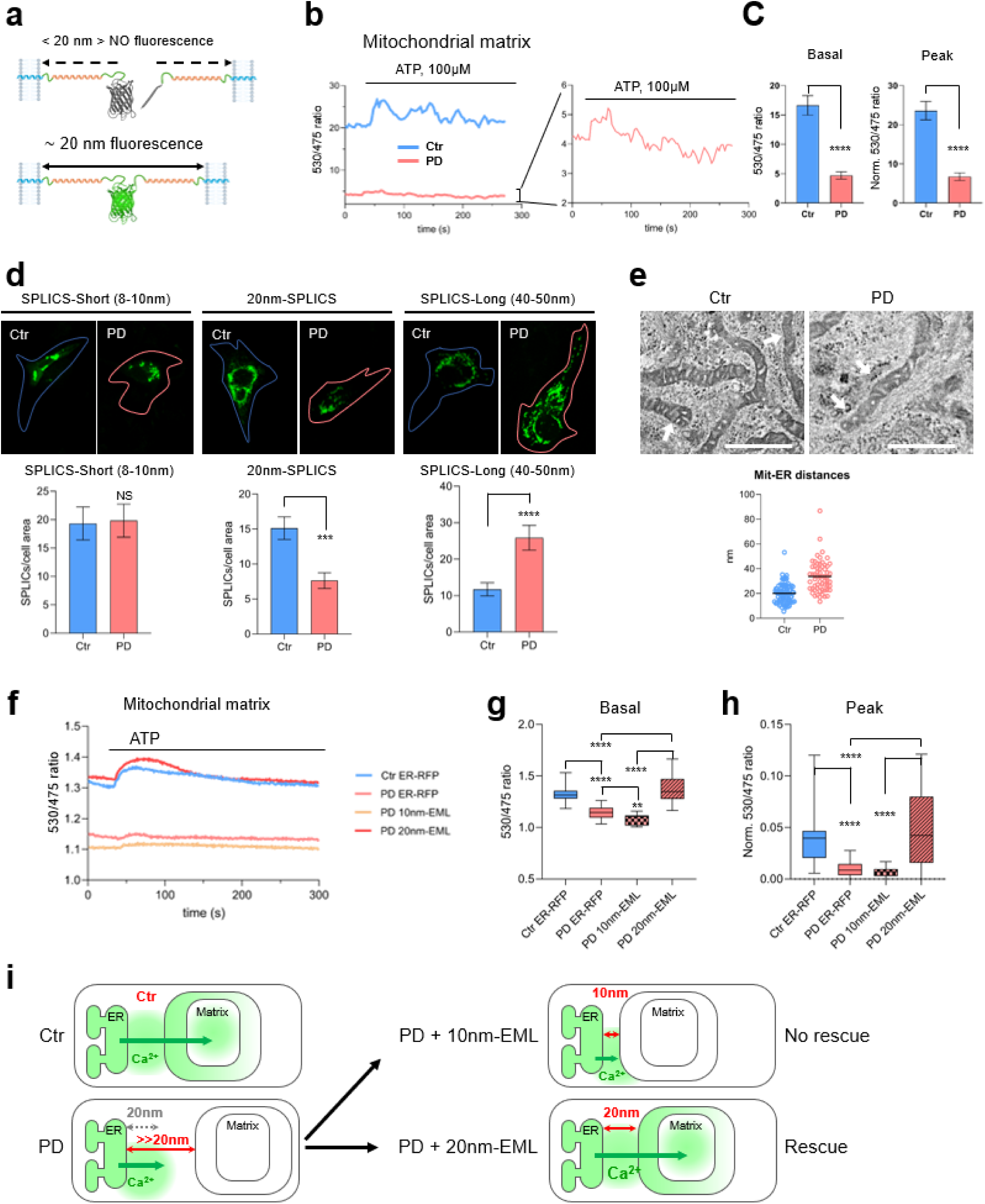
20nm ER-mitochondrial distance and mitochondrial Ca^2+^ are disrupted in astrocytes from Parkinson’s disease patients. (**a**) Design of 20 nm split-GFP contact site sensor (20nm-SPLICS). Representative traces (**b**) and quantifications (**c**) of ATP (100 µM)-induced Ca^2+^ transient in hiA from healthy subjects (Ctr) and from LRRK2^G2019S^ patients (PD). Data are expressed as mean ± SEM of basal 530/475 ratio (Basal) and peak of the response (Peak) from 56-63 cells from three independent experiments. **(d)** representative images and quantification of fluorescence SPLICS-Short, 20nm-SPLICS and SPLICS-Long transfected in hiA from healthy subjects (Ctr) and from LRRK2^G2019S^ patients (PD). Data are expressed as mean ± SEM of GFP area normalized to cell area from n = 19-22 cells from at least three independent transfections. Unpaired two-tailed Student’s t-test, *** p-value < 0.001; **** p-value < 0.0001. (**e**) Representative TEM micrographs and quantifications of the ER-OMM distance of hiA from healthy subjects (Ctr) and from LRRK2^G2019S^ patients (PD). Arrows indicate ER membrane adjacent to mitochondria. Scale bar = 500 nm. Data are mean ± SEM from 72-88 individual contact sites from 28 (Ctr) or 20 (PD) slices per genotype. Unpaired two-tailed Student’s t-test, **** p-value < 0.0001. Representative traces (**f**) and quantification (**g, h**) of Ca^2+^ responses in the mitochondrial matrix of Ctr-hiA and PD-hiA transfected with ER-mRFP, 10nm-EML and 20nm-EML. Box plot from 44-54 cells from three independent experiments. One-way ANOVA with Tukey post hoc test, **** p-value < 0.0001. (**i**) Cartoon illustrating MERCS alterations in hiA from LRRK2^G2019S^ patients (PD) compared with healthy subjects (Ctr). Enlargement of ER-OMM distance with specific reduction of 20 nm MERCS leads to failure of high [Ca^2+^] hot spot to reach mitochondria. The alterations are rescued by the expression of 20nm-EML but not by 10nm-EML.

As a model of pathological condition that explores the potential alteration of MERCS-mediated Ca^2+^ transfer, we capitalized on our recent characterization of hiPSC-differentiated astrocytes (hiA) from human subjects bearing a Parkinson’s disease (PD)-causing LRRK^G2019S^ mutation ^40^ (referred as to PD-hiA) ^41^. PD-hiA displayed no differences in basal or evoked cytosolic Ca^2+^ signals, while presenting severe defects of mitochondrial respiratory activity ^41^. Therefore, PD-hiA are suitable to explore if defects of mitochondrial respiration are caused by the deficiency of the ER-mitochondrial Ca^2+^ transfer and distance. We employed 4mtD3cpv probe to demonstrate that [Ca^2+^]_M_ signals were dramatically reduced in PD-hiA compared with control cells prepared from aged-pared non-PD subjects (Ctr-hiA) (Fig. 5c), suggesting impaired ER-mitochondrial Ca^2+^ transfer. To assess ER-mitochondria interaction, we used three available SPLICS sensors: SPLICS-Short, detecting distances at 8-10 nm; the newly-generated 20nm-SPLICS and SPLICS-Long, detecting contact sites at 40-50nm ^38,39^. As shown in Fig. 5d, no differences were found in MERCS at 8-10 nm distance (SPLICS-Short) between Ctr-hiA and PD-hiA; however, a dramatic decrease was observed in PD-hiA in the abundance of contact sites detected by 20nm-SPLICS. Instead, there was a significant increase of contact sites detected by SPLICS-Long in PD-hiA compared with Ctr-hiA (Fig. 5d), suggesting an enlargement of the distance between ER and mitochondria. To validate SPLICS data we quantified ER-mitochondrial distance on TEM images in a range from 5 to 100 nm. As shown in Fig. 5e, the average distance between ER and mitochondria increased from ∼20 to ∼38 nm, corroborating SPLICS quantification.

In order to investigate if mitochondrial Ca^2+^ uptake in PD-hiA could be rescued by stabilizing the ER-mitochondrial interaction at the 20 nm distance, optimal for Ca^2+^ transfer, we expressed control (ER-mRFP), 10nm-EML and 20nm-EML together with 4mtD3cpv probe in PD-hiA and stimulated cells with ATP. As shown in Fig. 5f-h, 10nm-EML did not rescue mitochondrial Ca^2+^ uptake, moreover, both basal Ca^2+^ level in the mitochondrial matrix and the ATP-induced Ca^2+^ transient in 10nm-EML-expressing PD-hiA were lower than in ER-mRFP-expressing PD-hiA cells. Strikingly, 20nm-EML overexpression fully rescued both the basal Ca^2+^ level in the matrix and the Ca^2+^ transient which were similar to those observed in Ctr-hiA expressing control ER-mRFP construct (Fig. 5g-h).. These results i) validate the importance of Ca^2+^ transfer at 20 nm MERCS suggesting that, contrarily to Alzheimer’s disease astrocytes, in which a shortening of the ER-mitochondrial distance is found ^15,42,43^ in PD-hiA bearing LRRK^G2019S^ mutation, impairment of mitochondrial bioenergetics ^41^ is due to enlargement of ER-mitochondrial distance, loss of 20 nm MERCS and inability of Ca^2+^, released from the ER, to reach mitochondria (Fig. 5i); ii) the defect of the mitochondrial Ca^2+^ uptake can be efficiently rescued by stabilizing the membranes at the distance of 20 nm by not 10 nm.

## DISCUSSION

Since the discovery of the close apposition between ER and mitochondrial membranes to warrant a low-affinity mitochondrial Ca^2+^ uptake ^8,9^, attempts have been made over the last three decades to estimate how ER-mitochondria interaction affects Ca^2+^ transfer. A range of ER-mitochondria distances between 10 and 25 nm has been hypothesized based on theoretical considerations ^14,15^. A more narrow range of 10-15 nm was considered for design of ER-mitochondria linking probes to quantify [Ca^2+^] in the ER-OMM cleft during IP_3_-mediated Ca^2+^ release ^12^. However, our results do not support this scenario, suggesting that a larger, ∼20 nm space is required to maximize ER-mitochondrial Ca^2+^ transfer. Although, at 10 nm, IP_3_Rs may be recruited into MERCS, the Ca^2+^ transfer is significantly less efficient compared with control, suggesting that at 10 nm the formation of functional IP_3_R-VDAC1 Ca^2+^ transferring units is inhibited. This conclusion is supported by the reduction of [Ca^2+^]_CL_ and [Ca^2+^]_M_ Ca^2+^ transients in 10nm-EML-expressing cells in spite of significant increase of the length of ER-OMM interface. Currently, it is unclear how 20 nm MERCS are organized and what is the mechanism of relocation of IP_3_Rs to MERCS. IP_3_Rs have several dozens of protein partners, many of which have been shown to promote IP_3_R-mediated Ca^2+^ signals at MERCS. Plausible candidates may include Sigma-1 receptor ^44^ and IRE1α ^45^ in the ER membrane, and Tom70 in the OMM ^46^. A hypothetical ∼7-10 nm gap between IP_3_R and VDAC might accommodate bridging proteins such as GRP75, suggesting a stoichiometric formation of IP_3_R-GRP75-VDAC complex ^26^. However, even if we show that both IP_3_R and VDAC1/3 are required for ER-mitochondrial Ca^2+^ transfer, our data do not support the hypothesis of physical interaction because we do not observe enrichment of both GRP75 and VDAC in 20 nm MERCS. Instead, our data are in line with a recent report suggesting that IP_3_R does not require physical linkage with a pore-forming protein in the OMM to warrant ER mitochondrial Ca^2+^ flux ^47^.

Impairment of mitochondrial oxidative metabolism and reduction of ATP production are common hallmarks of neurodegenerative diseases ^48^. Recently, we reported that in both AD and PD astrocytes, basal respiration, respiratory reserve capacity, and ATP synthesis are impaired ^41,42^. In AD astrocytes, bioenergetic deficit was associated with a reduction of mitochondrial Ca^2+^ uptake due to the shortening of the distance between ER and mitochondria with the specific increase of the interaction at 8-10 nm ^42,43^, i.e., the distance at which Ca^2+^ transfer is impaired, according to our current data. This prompted us to investigate if PD astrocytes had similar alterations of the ER-mitochondrial interaction found in AD astrocytes. Strikingly, and contrarily to AD astrocytes, we found a dramatic enlargement of the space between organelles with a reduction of 20 nm contacts associated with severe impairment of mitochondrial Ca^2+^ uptake. This finding suggests that, in pathological conditions, impairment of mitochondrial bioenergetics, as a consequence of the deficit of Ca^2+^ signals, may result from both shortening (in the case of AD) and enlargement (in the case of PD) of the distance between ER and mitochondria. Therefore, normalization of the ER-mitochondrial Ca^2+^ transfer through modulation of the ER-mitochondrial interaction may represent a universal therapeutic approach to normalize cellular homeostasis in different pathological conditions characterized by the impairment of mitochondrial function.

Taken together, in this study we demonstrate that the ER-OMM distance is a critical parameter for ER-mitochondrial Ca^2+^ transfer and OXPHOS. We show that a narrow range of distances close to 20 nm is optimal for Ca^2+^ flux between the organelles; the shortening of the distance below 20 nm dramatically affects both mitochondrial Ca^2+^ uptake and OXPHOS. We provide a framework and extend molecular tool for manipulating mitochondrial function through precise control over ER-mitochondrial distance.

## ONLINE METHODS

### DNA constructs

#### ER-mitochondrial linkers

5nm-EML and 10nm-EML were a kind gift from Georgy Hajnoszky, Jefferson University, USA) ^12,16^. Sequences of other linkers are reported in fig. S1:

Linkers are composed of targeting sequences for OMM (mouse AKAP1, residues 34-63) and ER (yeast Ucb6, residues 233-250) ^16^; flexible linkers, rigid α-helical spacers (yellow), and mRFP1 sequence (https://www.fpbase.org/protein/mrfp1/). EMLs were synthetized and cloned in pcDNA3.1 using Nhe1 and Xho1 restriction sites by GenScript (https://www.genscript.com/). EBFP-containing EMLs were synthetized by substituting mRFP1 sequence with EBFP2 sequence (https://www.fpbase.org/protein/ebfp2/).

#### Split-GFP contact site sensors (SPLICS)

Generation of SPLICS-Short and SPLICS-Long was described elsewhere ^38,39^. 20nm-SPLICS was synthetized by GenScript (https://www.genscript.com/) and cloned in pcDNA3.1. Sequence of 20nm-SPLICS is reported in fig. S9.

### Cell lines

HeLa (https://www.atcc.org/products/ccl-2), Huh-7 (https://huh7.com/) cells, HeLa-IP_3_R1-EGFP cells ^27^, HeLa-IP_3_R-TKO ^29^, MEF-VDAC1,3-KO ^34^, and their respective WT lines were maintained in complete culture media containing Dulbecco’s modified Eagle’s medium (DMEM; Sigma-Aldrich, Cat. D5671) supplemented with 10% fetal bovine serum (FBS, Gibco, Cat. 10270), 2 mM L-glutamine (Sigma-Aldrich, Cat. G7513), and 1% penicillin/streptomycin solution (Sigma-Aldrich, Cat. P0781).

### Cell transfection

3×10^4^ cells/well were resuspended in 250 µl of complete DMEM and 250 µl of transfection mix, and plated onto 13 mm glass coverslips in 24-well plates. For the transfection mix, Lipofectamine 2000 (Thermo Fisher Scientific, Cat. 11668-019) and plasmid, in a ratio 1:1, were mixed in Optimem (Gibco, Cat. 11058-021); after 3 h, transfection medium was replaced with complete medium. After 48 h, cells were used for experiments.

### Cell viability assay

Crystal violet is a viability assay that discriminates between alive and dead cells in culture by employing a blue/violet dye exclusively binding to DNA and proteins in well-adherent, viable cells. HeLa and Huh-7 were seeded, respectively at a density of 7.5×10^3^ and 15×10^3^ cells/well, and transfected with 5nm-EML, 20nm-EML, and ER-RFP on 96-well plates. 48h post-transfection, media was removed, and cells were fixed in methanol at 4°C. After incubation for 10-20 min with 50 µl/well of 0.1% crystal violet, the dye was carefully removed, and each well was washed with phosphate-buffered saline solution (PBS). Then, plates were allowed to dry for 12 h, and crystal violet was solubilized in 50 µl/well of 30% acetic acid. Lastly, absorbance at 595 nm was measured using Victor^3^V 1420 multilabel counter (Perkin Elmer).

### Transmission electron microscopy

For transmission electron microscopy (TEM) analysis, following trypsinization, 1×10^6^ cells were centrifuged at 900 rpm for 5 min and then fixed with 2.5% glutaraldehyde in culture medium, for 2 h at room temperature. The pellet was then rinsed in PBS, post-fixed in 1% aqueous OsO_4_ for 2 h at room temperature, and rinsed in H_2_O. Cells were pre-embedded in 2% agarose in water, dehydrated in a graded acetone scale, and then embedded in epoxy resin (Electron Microscopy Sciences, EM-bed812). Ultrathin sections (60–80 nm) were cut on a Reichert OM-U3 ultramicrotome, collected on nickel grids, and then stained with uranyl acetate and lead citrate. The specimens were observed with a JEM 1200 EX II (JEOL, Peabody, MA, USA) electron microscope operating at 100 kV and equipped with a MegaView G2 CCD camera (Olympus OSIS, Tokyo, Japan) ^49^. Images were analyzed via ImageJ 1.54F.

### Western Blot

48 h post-transfection, cells were lysed with lysis buffer (50 mM Tris-HCl (pH 7.4), sodium dodecyl sulphate (SDS) 0.5%, 5 mM EDTA, complemented with protease inhibitors cocktail (PIC, Millipore, Cat. 539133) and phosphatase inhibitor cocktail (Thermo Fisher Scientific, Cat. 78428), and collected in a 1.5 ml tube. Lysates were quantified with QuantiPro BCA Assay Kit (Sigma, Cat. SLBF3463). According to the relative abundance of the protein of interest, 20-40 µg of proteins were mixed with the right amount of Laemmli Sample Buffer 4X (Bio-Rad) and boiled. Then samples were loaded on a 6-12% polyacrylamide-sodium dodecyl sulphate gel for SDS-PAGE. Proteins were transferred onto nitrocellulose membrane, using Mini Transfer Packs or Midi Transfer Packs, with Trans-Blot® Turbo ^TM^ (Bio-Rad) according to the manufacturer’s instructions (Bio-Rad). The membranes were blocked in 5% skim milk (Sigma, Cat. 70166) for 45 min at room temperature. Subsequently, membranes were incubated with the indicated primary antibody overnight at 4°C. Anti-β-Actin was used to normalize protein loading. A list of primary antibodies is provided in Supplementary Table 1.

Goat anti-mouse IgG (H+L) horseradish peroxidase-conjugated (Bio-Rad, 1:5000; Cat. 170-6516,) and Goat anti-rabbit IgG (H+L) horseradish peroxidase-conjugated secondary antibodies (Bio-Rad, 1:5000; Cat. 170-6515,) were used. Detection was carried out with SuperSignal^TM^ West Pico/femto PLUS Chemiluminescent Substrate (Thermo Scientific, Cat. 34578), based on the chemiluminescence of luminol and developed using ChemiDoc^TM^ Imaging System (Bio-Rad).

### Time-lapse ratiometric fluorescent imaging

Imaging of Fura-2, GAP3, 4mtD3cpv, and ROMO-GemGeCO Ca^2+^ probes was performed using an epifluorescent Leica DMI6000B microscope equipped with an S Fluor 40×/1.3 objective, a Polychrome V monochromator (Till Photonics, Munich, Germany), a Photometrics DV2 dual-imager (Teledyne Photometrics, Tucson, US). For imaging of mitochondria, an internal lens with a 1.6 optical increment was used. Images were acquired by a Hamamatsu cooled CCD camera (Hamamatsu Photonics, Hamamatsu City, Japan) and registered using MetaFluor software (Molecular Devices, Sunnyvale, CA, USA). Microsoft Excel and GraphPad Prism were used for offline analysis and figure preparation.

#### Mitochondrial Ca^2+^ imaging

Mitochondrial Ca^2+^ dynamics were monitored with 4mtD3cpv sensor, a genetically encoded Ca^2+^ indicator targeted to the mitochondrial matrix ^22^. 48h post-transfection, expression of 4mtD3cpv was checked and mitochondrial matrix calcium dynamics were monitored. Coverslips were washed with KRB solution (125 mM NaCl, 5 mM KCl, 1 mM Na_3_PO_4_, 1 mM MgSO_4_, 5.5 mM glucose, 20 mM HEPES, pH 7.4), transferred to an acquisition chamber, and mounted on the stage of the microscope. Samples were illuminated at 420 nm and simultaneously acquired at 475 nm (donor, ECFP) and 530 nm (acceptor, circularly permuted (cp) Venus). CpVenus/ECFP ratio was calculated online using MetaFluor software. After acquisition of basal Ca^2+^ levels (first 30 s of acquisition), the cells were stimulated with 100 μM ATP. Regions of interest (ROIs) were defined around individual mitochondria.

#### Fura-2 Ca^2+^ imaging

Cells were plated onto 24 mm round coverslips (3×10^4^ cell/coverslip), and loaded with 2.5 μM Fura-2/AM (Cat. No. F1201, Life Technologies, Milan, Italy) in the presence of 0.005% Pluronic F-127 (Cat. No. P6867, Life Technologies) and 10 μM sulfinpyrazone (Cat. S9509, Sigma) in KRB solution. After loading (30 min in the dark at RT), cells were washed once with KRB solution and allowed to de-esterify for 30 min. After this, the coverslips were mounted in an acquisition chamber and placed on the stage of the microscope, and cells were alternately excited at 340 and 380 nm; the fluorescent signal was collected through a 510/20 nm bandpass filter. The cells were stimulated with 100 μM ATP. For comparison of Ca^2+^ dynamics, measured as an amplitude of Ca^2+^ increase from the baseline level, Fura-2 ratio values were normalized using the formula (Fi-F0)/F0 [referred to as Normalized (Norm.) Fura Ratio].

#### Endoplasmic reticulum Ca^2+^ imaging

ER Ca^2+^ dynamics were monitored with ER-GAP3, a genetically encoded Ca^2+^ sensor, targeted to the ER lumen (referred to as GAP3) ^24^. 48 h post-transfection, expression of GAP3 was checked and ER calcium dynamics were monitored. Coverslips were mounted in a chamber in KRB solution and placed on the stage of the microscope. Cells were alternately excited at 405 and 470 nm, and the fluorescent signal was acquired using a 510/20 nm bandpass filter. After recording basal signal for 30 s, KRB solution was removed and replaced with a Ca^2+^-free solution (KRB + 500 µM EGTA). After allowing the signal to stabilize for additional 30 s, cells were stimulated with 100 µM ATP and 100 µM Tert-butylhydroquinone (TBHQ), and the response was recorded for 300 s.

#### Ca^2+^ imaging in mitochondrial cristae lumen

Ca^2+^ dynamics in the mitochondrial cristae lumen (CL) were monitored with ROMO-GemGeCO, a genetically encoded Ca^2+^ indicator localized to the cristae lumen space ^25^. 48h post-transfection, coverslips were washed with KRB solution and transferred to the acquisition chamber and mounted on the stage of the microscope. Samples were illuminated at 420 nm and simultaneously acquired at 475 nm and 530 nm. 530/475 nm ratio was calculated online using MetaFluor software. After acquisition of basal Ca^2+^ levels (first 30 s of acquisition), the cells were stimulated with 100 μM ATP. Regions of interest (ROIs) were defined around individual mitochondria.

### Immunofluorescence

Cells, transfected or not according to the experimental design, were grown on 13 mm glass coverslips, fixed with 4% formaldehyde, permeabilized (7 min in 0.1% Triton X-100 in PBS), blocked in 1% gelatin, and immunoprobed with an appropriate primary antibody (diluted in PBS supplemented with 1% gelatin) overnight at 4°C. After 3 times washing in PBS, an Alexa-conjugated secondary antibody (1:300 in PBS supplemented with 1% gelatin) was applied for 1 h at room temperature (RT). The following primary antibodies were used: anti-IP_3_R (rabbit, 1:500, Abcam, Cat. AB108517) and anti-VDAC1/3 (mouse, 1:250, Abcam, Cat. AB14734). Secondary antibodies were as follows: Alexa Fluor 488 anti-mouse IgG, Alexa Fluor 555 anti-rabbit IgG (all secondary antibodies were from Molecular Probes, Life Technologies, Monza, Italy). Nuclei were counter-stained with 4′,6-diamidino-2-phenylindole (DAPI). Images were acquired by Zeiss 710 confocal laser scanning microscope equipped with EC Plan-Neofluar 40×/1.30 Oil DIC M27 objective and Zen software or with a Leica SP8 LSCM equipped with a white light laser, and HCX PL APO 40X/1.25-075 OIL CL objective and LAS X software. Image stacks were processed offline to major intensity projections. Thresholding and area calculations were done using Fiji package of Image J software v.1.52p. Data are expressed as fluorescence intensity/cell area ratio. For measuring colocalization of two fluorophores we used an ImageJ JACoP toolbox for subcellular colocalization analysis ^50^.

### Proximity ligation assay (PLA)

1.75 x 10^4^ cells/well were plated in 8 wells chamber (IBIDI, Cat: 80806) and transfected with ER-mRFP, 5nm-EML, 10nm-EML and 20nm-EML. After 40h, PLA was performed according to manufacturer’s instructions (Duolink® Proximity Ligation Assay, Sigma). Briefly, cells were fixed in 4% paraformaldehyde, and incubated with primary antibodies anti-IP_3_R (1:500) and anti-VDAC1/3 (1:100) for 16 h at 4°C. Duolink PLA probe incubation, according to the primary antibody species, was carried on for 1 h at 37°C, and then the development of the signal was obtained by Duolink green fluorescence detection reagent by ligation and amplification reactions. Duolink in situ mounting media with DAPI was used. Images were acquired by Zeiss 710 laser scanning confocal microscope (LSCM) equipped with EC Plan-Neofluar 40×/1.30 Oil DIC M27 objective and Zen software. Images were acquired under non-saturating conditions and analyzed with Fiji ImageJ 1.52p software. Fluorescence was measured for the entire cell area (CTCF) = Integrated Density — (Area of selected cell X Mean background fluorescence).

### Generation of stable lines expressing ER-mRFP and 20nm-EML

To isolate MERCS-enriched cellular fraction, stable HeLa cell lines expressing ER-mRFP and 20nm-EML were generated. To generate ER-mRFP and 20nm-EML-expressing lentiviral backbones, ORFs were excised from pcDNA3.1 using Nhe1 and Xho1 restriction sites and cloned into Xba1 and Sal1 restriction sites, respectively, of pRRLSIN.cPPT.hCMV-GFP.WPRE vector. Lentiviral particles were produced and concentrated using ultracentrifugation protocol described elsewhere ^51^. For generation of stable lines, 24 h after plating (10^4^ cell/well in 24-well plates), HeLa cells were infected with lentiviral particles expressing ER-mRFP and 20nm-EML at MOI from 5 to 20. The wells presenting more than 50% of infected cells, as detected by fluorescence of reporter proteins, were further processed. Cells were expanded, and ER-mRFP and 20nm-EML expressing cells were enriched using fluorescence-activated cell sorting (S3e Cell Sorter, Bio-Rad, Segrate, Milano). Sorted cells were expanded, cryopreserved and stored in liquid nitrogen until needed.

### Isolation of MERCS

HeLa stably expressing ER-mRFP (also referred to as ER-RFP) and 20nm-EML were plated at a concentration of 0.5 x10^6^ cells/dish in 10 cm Petri dishes (50 dishes per line). 48 h later, cells were washed twice with PBS, detached with trypsin, and then collected in 50 ml centrifuge tubes. Cells were pelleted, and the pellets were processed according to the protocol described elsewhere ^52^, using an Eppendorf CR30NX ultracentrifuge equipped with an R25ST rotor.

### ATPlite™ assay

Cells were seeded, respectively, at a density of 7.5×10^3^ and 15×10^3^ cell/well, and transfected with 5nm-EML, 20nm-EML, and ER-mRFP on 96-well plates. 48h post-transfection, 30 µl/well of mammalian cell lysis solution were added, and lysis was favored by shaking the plate at 400/500 rpm for 5 min. Then, 30 µl/well of substrate buffer solution (containing Luciferase and D-Luciferin) were added, and the plate was put again at 400/500 rpm for 5 min, protected from light according to manufacturer’s instructions (PerkinElmer, Cat. ATPLT-0415). After another 10 min of incubation, luminescence was measured using Victor^3^V 1420 multilabel counter (Perkin Elmer).

### Mitochondrial membrane potential determination

JC-1 dye (Cayman, Ann Arbor, Michigan, USA, Cat.15003) was used according to manufacturer’s instructions to assess mitochondrial membrane potential (ΔΨm). Control and EMLs expressing cells were resuspended in complete media to a final cell density of 1×10^6^ cells. JC-1 dye was added to the cell suspension at a final concentration of 1 µg/µl and incubated for 15 min at 37°C and 5% CO_2_. A negative control sample was prepared by adding an equal volume of vehicle to the cell suspension. After washing with warm PBS, cells were resuspended in PBS + EDTA (500 µM), and immediately acquired with a flow cytometer (Accuri C6 Plus BD 660517). The fluorescence was measured with excitation wavelength at 485 nm, dual emission filters at 529 and 590 nm, and cut-off at 515 nm. 10^5^ events were acquired for each sample using the following parameters: forward scatter (FSC), side scatter (SSC), PE (red, JC-1 aggregate), and FITC (green, JC-1 monomer) fluorescence channels. Gating strategies were applied to exclude cell debris and doublets based on FSC and SSC properties. The mitochondrial membrane potential was determined as a ratio of JC-1 aggregates (red fluorescence) to monomers (green fluorescence). As a positive control, cells were treated with FCCP 10 µM (Tocris, Cat. 0453), for 5 min. Data analysis was performed using FlowJo software v10. Gated events were plotted on a bivariate dot plot to analyze the distribution of JC-1 fluorescence.

### Oroboros high-resolution respirometry

Cellular respiration rates in real-time of intact control and EMLs expressing cells were measured using an Oroboros oxygraph-2K high-resolution respirometer (Oroboros Instruments, Innsbruck, Austria). The "substrate, uncoupler, inhibitor, titration" (SUIT) protocol, specifically SUIT-003_O2_ce_D012, was employed following the guidelines recommended by the manufacturer. Cells were detached from the plate, using trypsin-EDTA (Gibco, Cat 25200056), counted, and resuspended in pre-warmed respiration medium MiR05 (0.5 mM EGTA, 3.0 mM MgCl_2_, 60 mM potassium lactobionate, 20 mM taurine, 10 mM KH_2_PO_4_, 20 mM HEPES, 110 mM sucrose, 1 g/L bovine serum albumin, pH 7.1) to achieve a final cell density of 1×10^5^ cells/ml. Pairs of ER-mRFP + 10nm-EML or ER-mRFP + 20nm-EML-expressing cells were analyzed simultaneously in the neighboring chambers, and oxygen concentration and flux were recorded using DatLab software (Oroboros). Baseline oxygen consumption rates were identified for each chamber during the “Routine” phase in the presence or absence of pyruvate (5 mM) stimulation, used to sustain TCA-linked respiration in MiR05 medium. Subsequently, oligomycin (5 nM) was added to inhibit ATP synthase and assess the uncoupled respiration (“Leak” phase). The protonophore carbonyl cyanide 4-(trifluoromethoxy) phenylhydrazone (FCCP) was then titrated (0.05 μM increments) until plateau of oxygen flux, indicative of maximal respiration, was achieved [“Electron transport (ET)” phase]. Finally, 1 μl each of rotenone (0.5 μM) and antimycin A (2.5 μM) were sequentially added to inhibit ETC complexes I and III, respectively, and to identify ET-independent respiration (ROX phase). Rates of O_2_ consumption (flux) were normalized to total protein content. Briefly, at the end of the experimental procedure, the cellular suspension from the two chambers was centrifuged at 1,000 × g for 5 min. The cellular pellets were lysed in 200 µL of lysis buffer (10 mM HEPES, 60 mM KCl, 1 mM EDTA, 0.075% NP40, 1 mM DTT) and then centrifuged at 15,000 × g for 15 min at 4 °C. Protein concentration in the supernatant was measured with Bradford Reagent (Sigma, Cat. B6916).

### Human induced astrocytes (hiA) derived from fibroblast of healthy donors and Parkinsońs disease patients with LRRK2^G2019S^ mutation

Human fibroblasts were obtained from two healthy donors (Ctrl1 was purchased from AXOL-#AX0019 and Ctrl 2 from the Coriell stem cell bank-#ND291914) and two Parkinsońs disease patients with LRRK2^G2019S^ mutation (PD1 from the Coriell stem cell bank-#PD33878, and PD 2 provided by the BioDonostia Hospital, San Sebastian, Spain). Generation and characterization of hiA have been described elsewhere ^41^. hiA were differentiated from embryoid bodies, obtained in Aggrewell 800 (STEMCELL) from human iPSCs. Differentiation of NPC to progenitor astrocytes was triggered using the astrocyte differentiation medium (STEMdiff astrocyte differentiation #100-0013, StemCell). To maintain the appropriate cell density (70% of confluence), cells were passed every week in the same coating mix (human laminins 50% LN211 and 50% LN111 from Biolamina) for 21 days. Finally, astrocyte progenitor cells were maturated in the Astrocyte Maturation Medium (STEMdiff astrocyte maturation #100-0016, StemCell) for 60 to 75 days before running the experiments.

### Statistical analysis

Statistical analysis was performed with GraphPad Prism software (Graphpad software Inc., La Jolla, CA). A two-tailed unpaired Student’s t-test was used to compare two samples. To compare three or more samples, one-way ANOVA was used, followed by Tukey post hoc test, unless otherwise specified. A p-value < 0.05 was considered statistically significant. A full report on statistical analysis is presented in Supplementary Tables S2 and S3.

