## Supplementary material for "ER-mitochondria distance is a critical parameter for efficient mitochondrial Ca^2+^ uptake and oxidative metabolism"

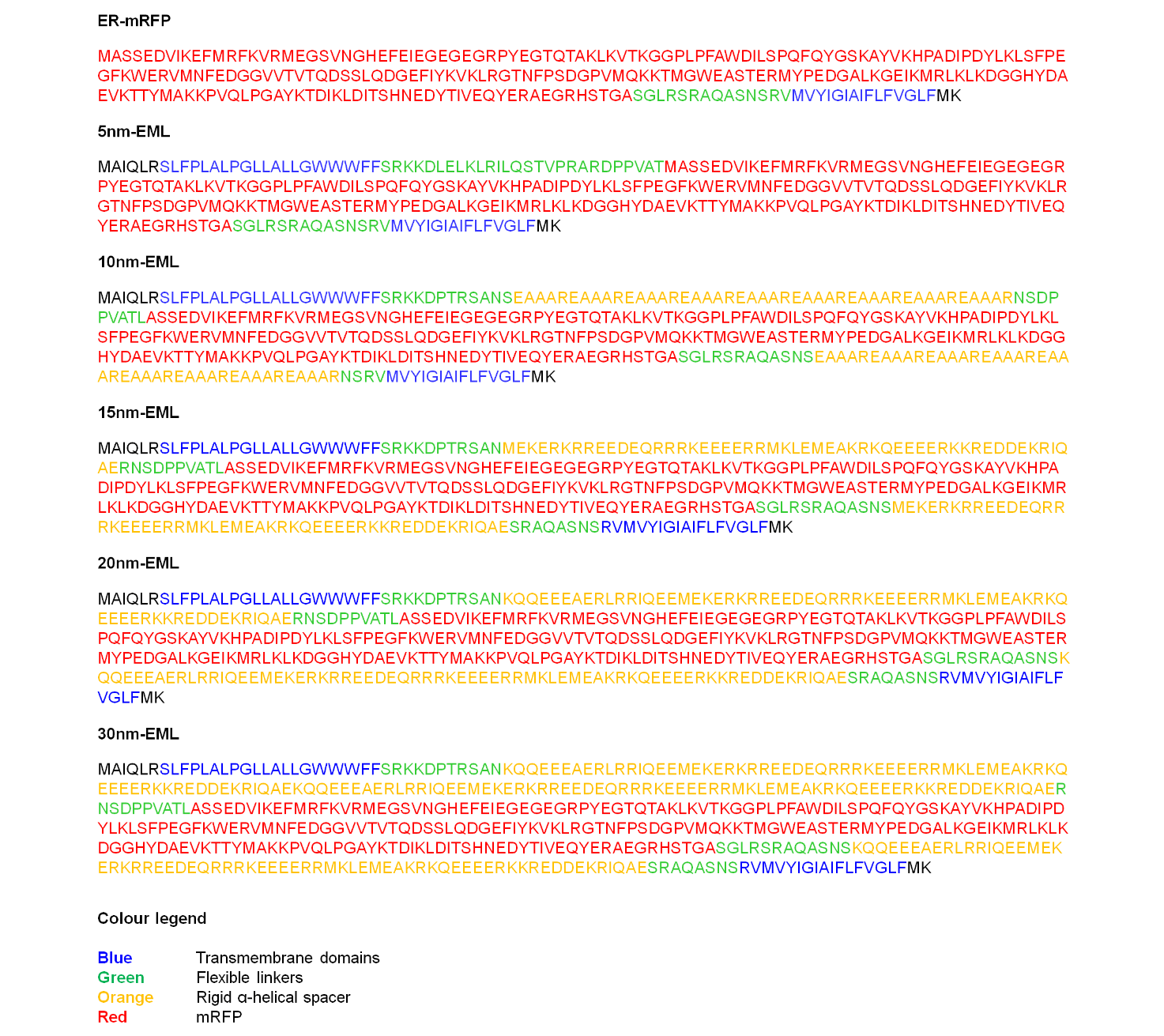

**Extended Fig. 1. Sequence of ER-mitochondrial linkers.** Color legend: **Blue**, Transmembrane domains; **Green**, flexible linkers; **Orange**, rigid α-helical spacer; **Red**, mRFP.

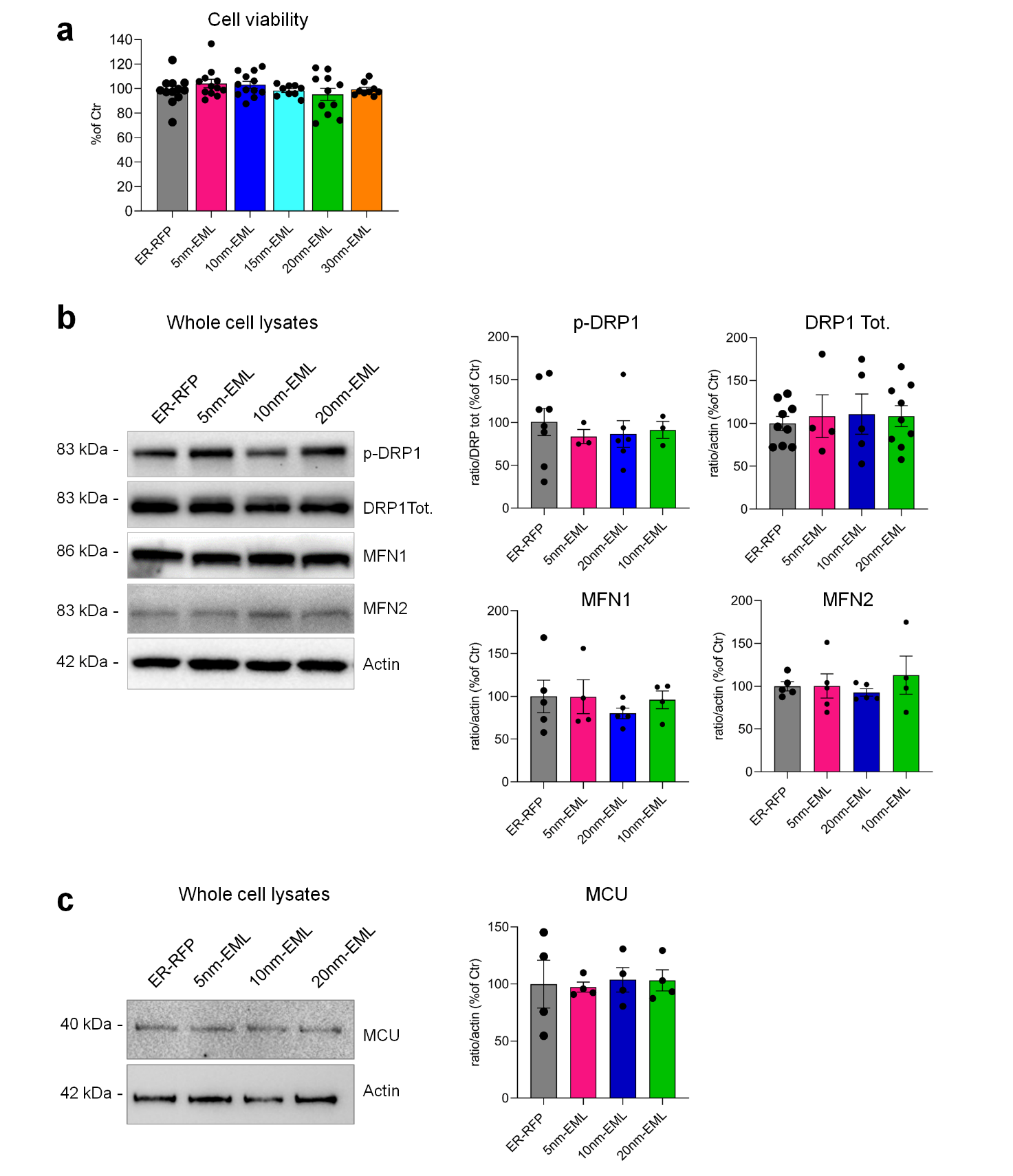

**Extended Fig. 2. Cell viability and expression of mitochondrial proteins in EMLs-expressing HeLa cells.** (**a**) cell viability determined using crystal violet assay. Data are mean ± SEM 595 nm absorbance from 3-4 independent experiments performed in triplicate. Representative images and quantifications of Western blots of proteins implicated in mitochondrial dynamics such as DRP1, p-DRP1, MFN1 and MFN2 (**b**) or Ca^2+^ uptake (MCU) (**c**). Data are mean ± SEM band intensity from at least 3 independent preparations. One-way ANOVA shows no statistically significant differences for all panels.

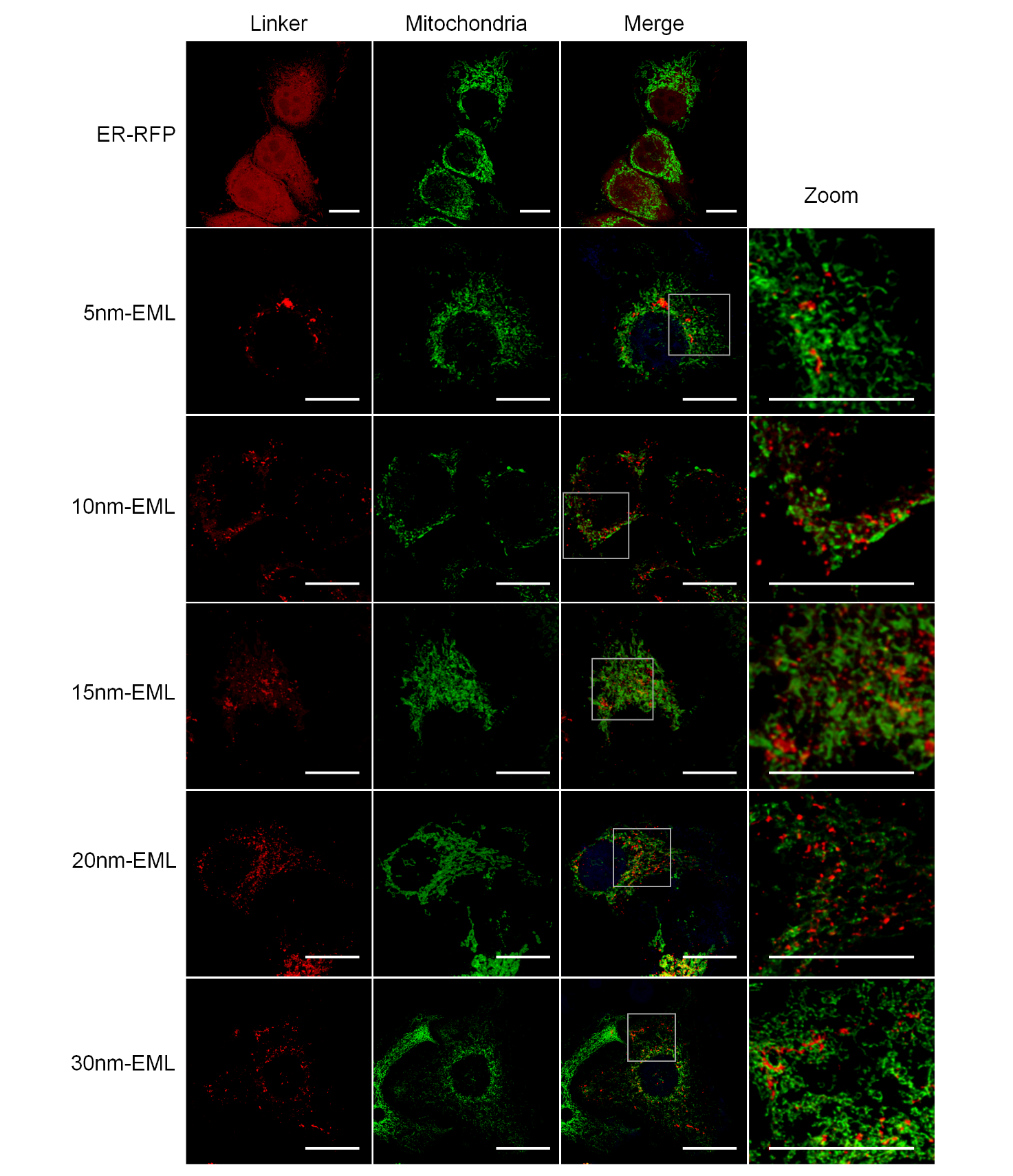

**Extended Fig. 3. Validation of EMLs.** Representative images show localization of EMLs (red, linker) and their colocalization with a mitochondrial marker (4mtD3cpv) (mitochondria). Rectangles in merged images are areas enlarged in Zoom images. Scale bars = 20 μm.

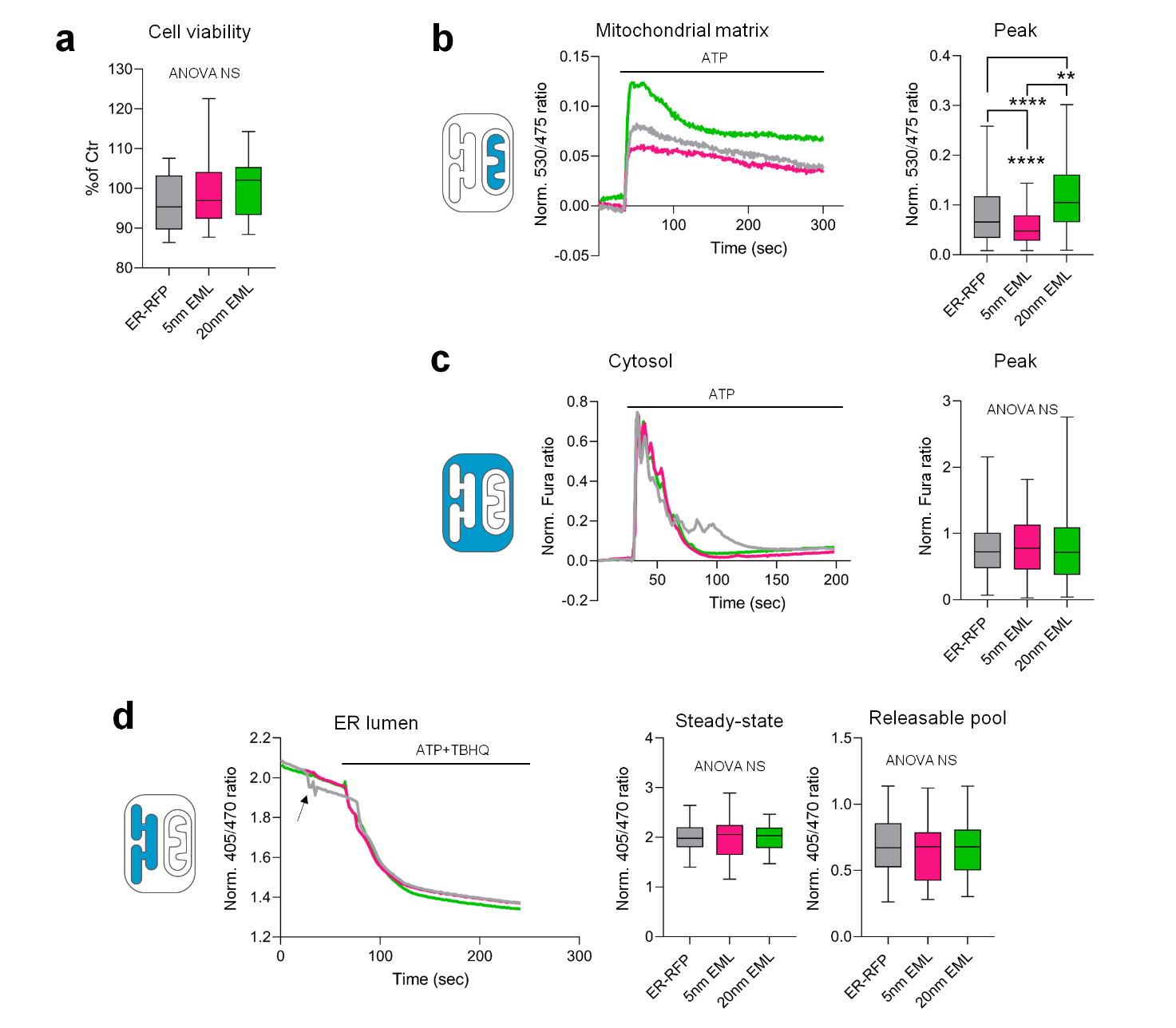

**Extended Fig. 4. Viability and Ca^2+^ handling in EML-expressing hepatocellular carcinoma Huh-7 cells**. (**a**) Whisker plot quantification of viable cells in EML-expressing Huh-7 cells performed using crystal violet assay. (**b**, **c** and **d**) representative traces and quantifications of Ca^2+^ signals in mitochondrial matrix (**b**), cytosol (**c**) and ER lumen (**d**) of EML-transfected Huh-7 cells. In (**a**) data are from four independent experiments performed in triplicate. One-way ANOVA non-significative. In (**b** and **c**) whisker plots show data from 106-275 cells collected from 12 coverslips from at least three independent experiment. (**c, d**) One-way ANOVA, Tukey post hoc test, ** p-value < 0.01; **** p-value < 0.0001. In (**d**) arrow indicates an artifact produced during medium exchange.

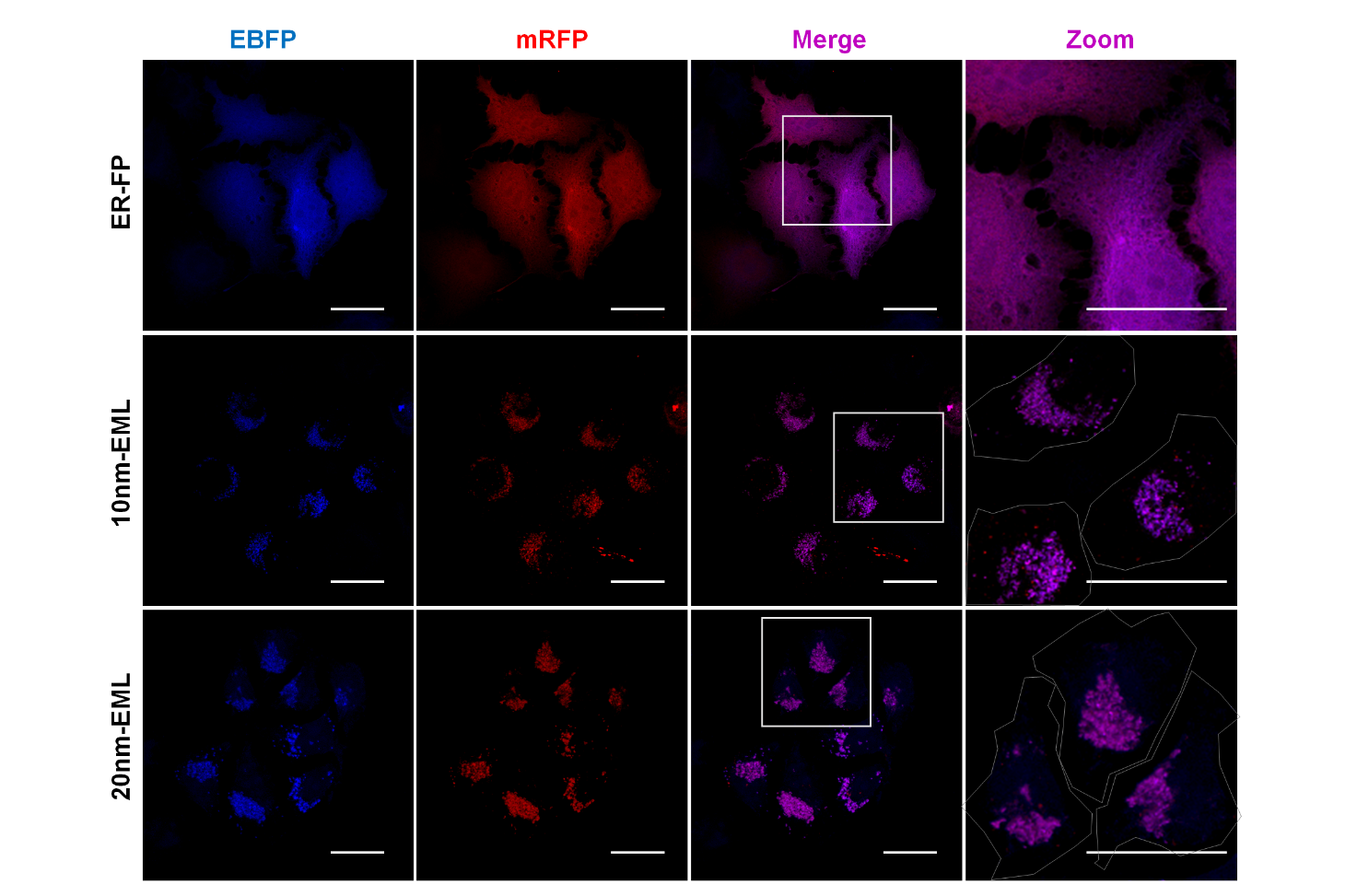

**Extended Fig. 5. Validation of blue EML variant expressing EBFP.** mRFP and EBFP-expressing ER-FP, 10nm-EML and 20nm-EML were co-expressed in HeLa cells. Note complete colocalization between red and blue EML variants. Rectangles on merged images indicate fields enlarged in Zoom images. Scale bar = 20 µm.

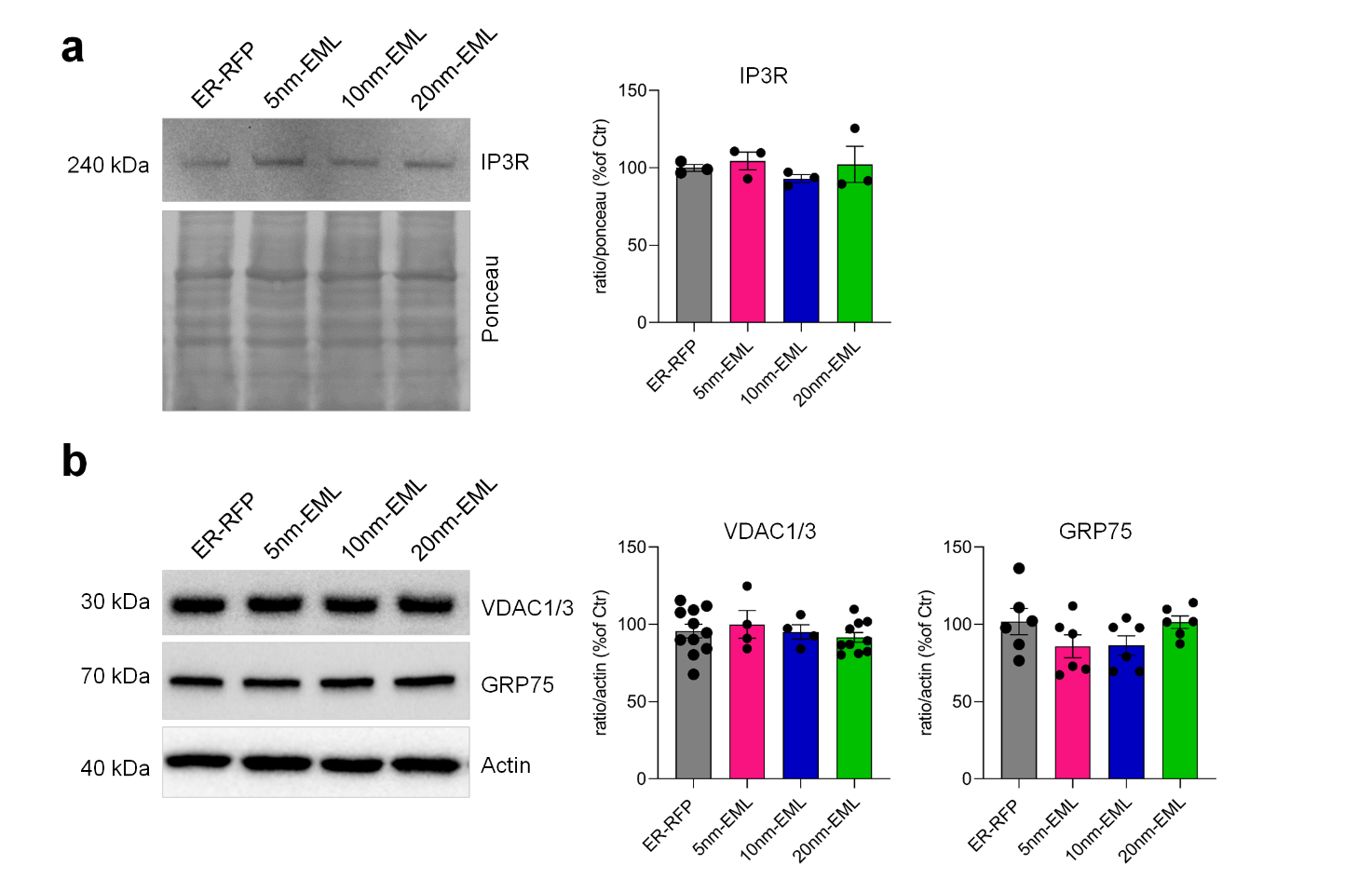

**Extended Fig. 6. Expression of IP_3_R, VDAC1/3 and GRP75 in total cell lysates.** Representative blots of total cell lysates and quantification of IP_3_R (**a**) and VDAC1/3 and GFP75 (**b**). Data are expressed as mean ± SEM of n = 3 (IP_3_R) or n = 4-11 (for VDAC1/3 and GFP75) experiments. One-way ANOVA shows no statistically significant differences.

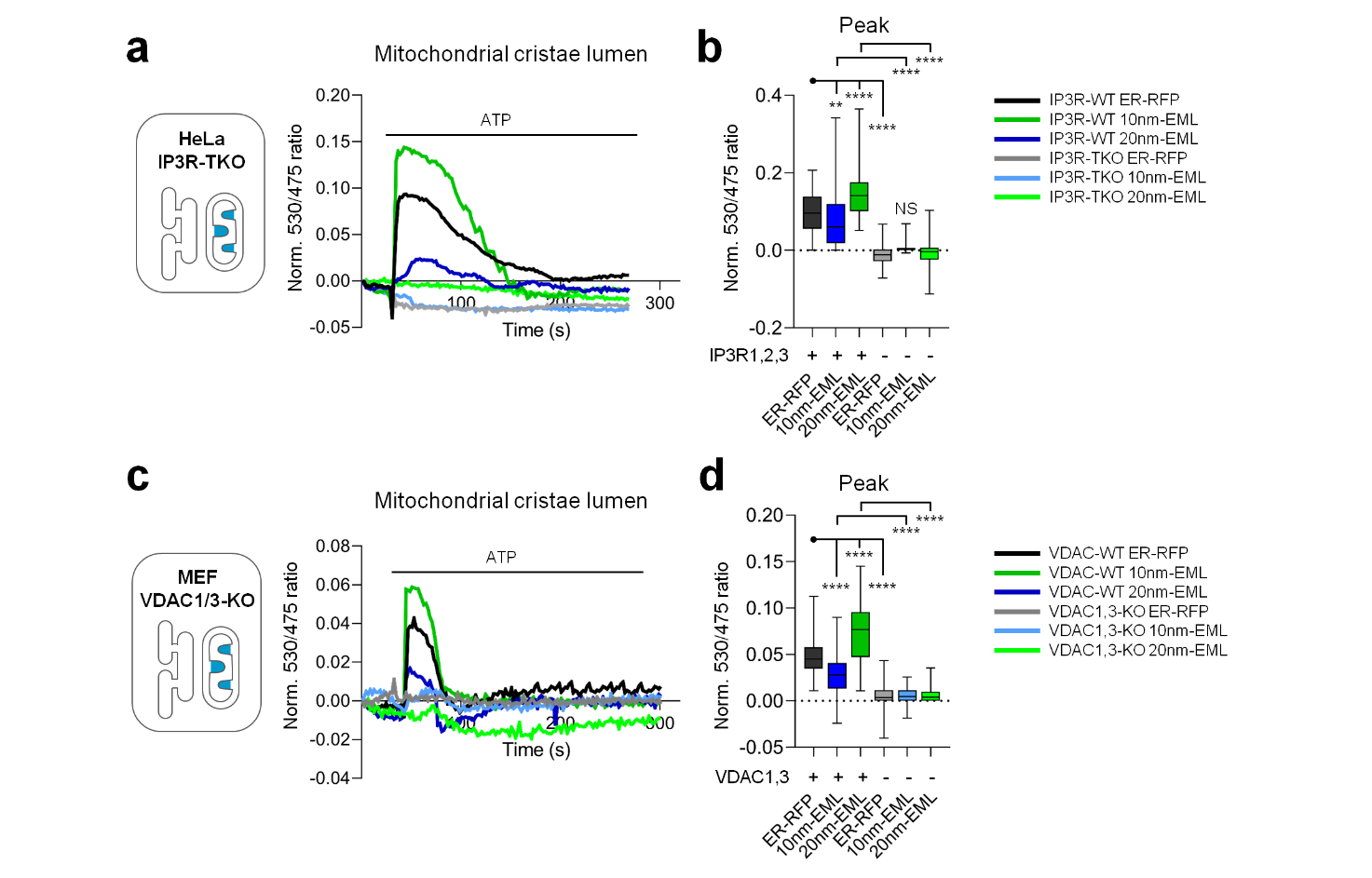

**Extended Fig. 7. IP_3_R and VDAC1/3-dependence of mitochondrial Ca^2+^ signals in EML-expressing cells.** Representative traces (**a, c**) and quantification of the amplitude of Ca^2+^ response in mitochondrial cristae (**a, d**) in IP_3_R-TKO (**a**, **b**) and VDAC1/3-KO cells (**c, d**) transfected with ER-RFP, 10nm-EML and 20nm-EML. Whisker plots report data from 59-102 cells from at least three independent experiments performed in triplicate. One-way ANOVA, Tukey post hoc test. ** p-value < 0.01; **** p-value < 0.0001.

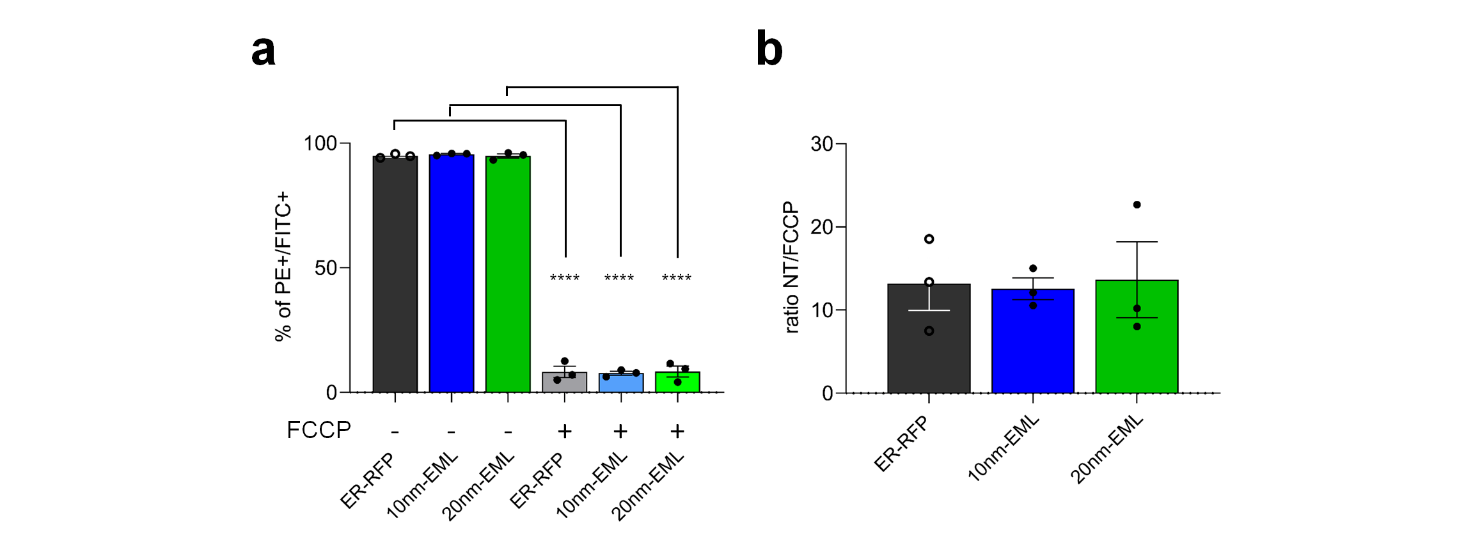

**Extended Fig. 8. Mitochondrial membrane potential (ΔΨm) is not affected by the expression of control (ER-EBFP), 10nm-EBFP or 20nm-EBFP EMLs**. (**a**) Quantification of G-aggregates/monomers ratio of JC-1 indicates no difference in ΔΨm between samples. FCCP (10 μM) was used as positive control. (**b**) Ratio between non-treated (NT) and FCCP (10 µM)-treated cells. Data are mean ± SEM of 10^5^ gated cells from three independent experiments. One-way ANOVA, Tukey post hoc test. **** p-value < 0.0001.

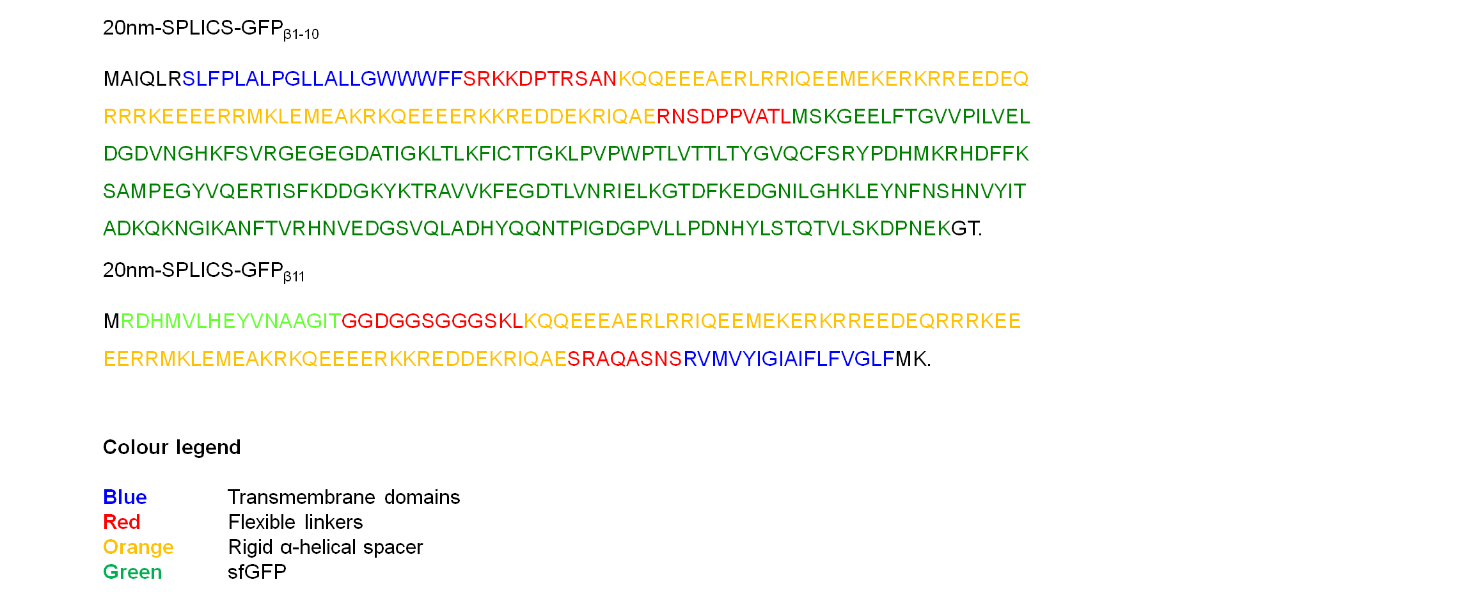

**Extended Fig. 9. Sequence of 20nm-SPLICS.** Color legend: **Blue**, transmembrane domains; **Red**, flexible linkers; **Orange**, rigid α-helical spacer; **Dark green**, superfolded (sf)GFPβ1-10; Light green, sfGFPβ11.

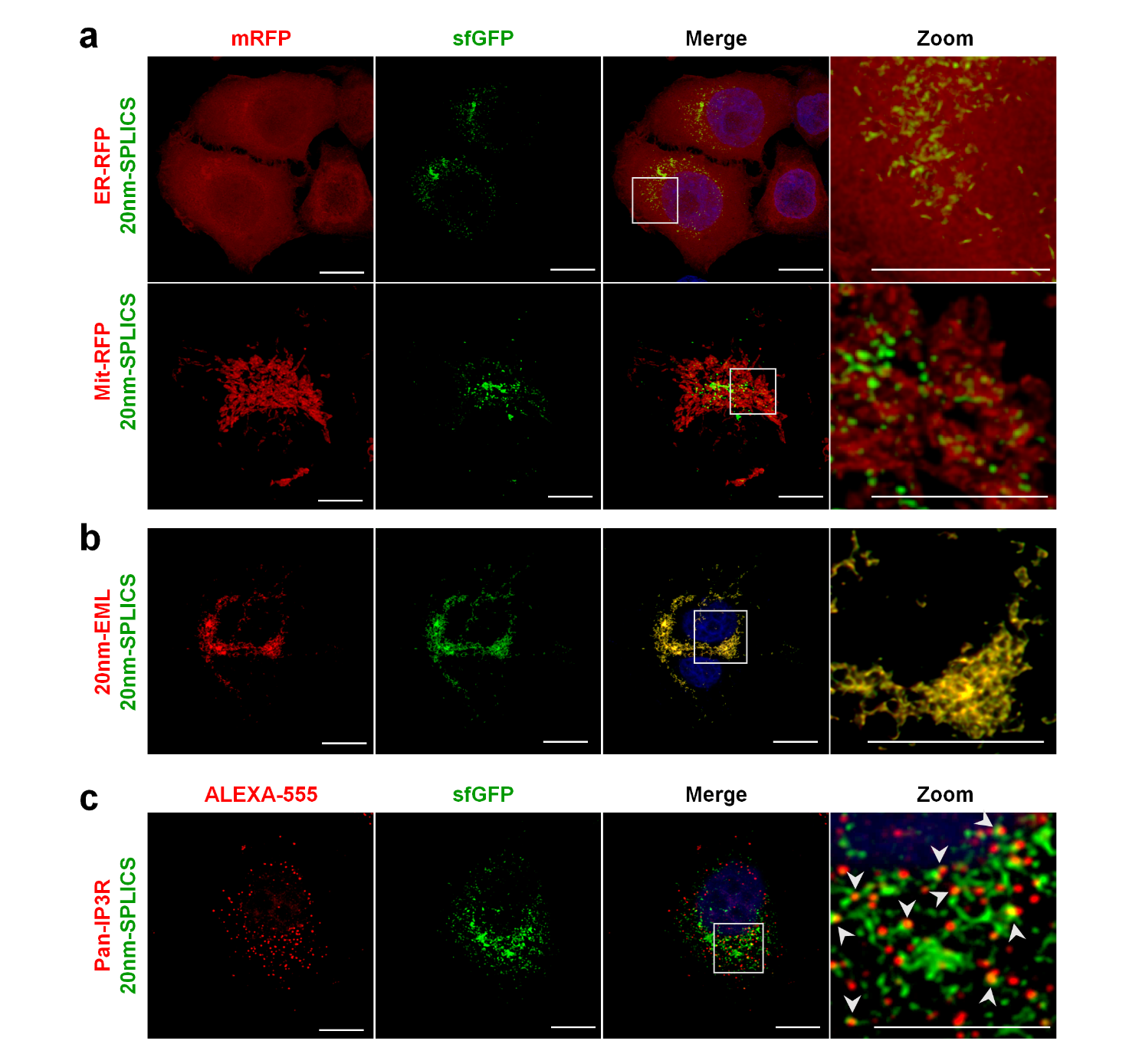

**Extended Fig. 10. Characterization of 20nm-SPLICS.** Colocalization of 20nm-SPLICS signal (sfGFP) with ER (ER-RFP) and mitochondria (Mit-RFP) (**a**) and with 20nm-EML (**b**). (**c**), Colocalization of 20nm-SPLICS signal (sfGFP) with IP_3_R detected using anti-pan-IP_3_R antibody (Alexa 555). Scale bar = 20 μm. White rectangles in merged images indicate area enlarged in Zoom images.
